# Spatial distribution of the proteome in human body and cancers

**DOI:** 10.1101/2025.02.14.638212

**Authors:** Liang Yue, Wenhao Jiang, Sainan Li, Meng Luo, Ning Fan, Xiaolu Zhan, Kexin Chen, Tian Lu, Fang Guo, Dongwei Li, Weigang Ge, Zongxiang Nie, Mengge Lyu, A Jun, Yingrui Wang, Yingdan Chen, Zhenhai Fu, Nan Xiang, Lu Li, Fengchao Yu, Guo Ci Teo, Alexey I Nesvizhskii, Meng Wang, Michael P. Snyder, Ben Collins, Ruedi Aebersold, Fei Xu, Yi Zhu, Tong Liu, Yan Li, Tiannan Guo

## Abstract

A comprehensive spatial distribution of the proteome in human body and cancers is fundamental for understanding human biology and diseases including cancers. Here, we present an anatomically resolved human proteome derived from 1781 benign and malignant samples from 58 major tissue types encompassing 251 specific tissues and 25 carcinomas. Based on a spectral library covering over 75% of the human protein-coding genes with 208 understudied and 82 missing proteins characterized, we quantified over 13,000 proteins in these samples using data-independent acquisition proteomics. This data resource presents the so far most comprehensive quantitative proteomic landscape of human tissues and common carcinomas. It allows systematic evaluation of tissue-specific drug responses, identification of drug candidates that may be repurposed as antineoplastics, and discovery of novel targets for anticancer therapy. This resource, available as an online knowledgebase, refines our knowledge of spatial distribution of the human proteome and tumor-specific protein modulation.

## INTRODUCTION

Grounded in a genetic blueprint, the development of diverse organs and tissues, each serving distinct functions, gives rise to the complex and sophisticated architecture of the human body. Elucidating the molecular intricacies underlying proteome variation across the diverse tissues and organs in healthy and pathological status is crucial for enhancing our understanding of human biology and the pathogenesis of diseases. Several transcriptomic repositories, such as ArrayExpress^1^, RNA-Seq Atlas^2^ and BioGPS portal^3^, have provided initial annotations for tissue expression. The Adult GTEx project has further enhanced our understanding by collecting genomic and transcriptomic data from numerous non-diseased tissue sites^4-6^.

Proteins are gears of life activities and major drug targets. A comprehensive analysis of proteins is anticipated to enhance our molecular understanding of complex tissues. The Human Protein Atlas (HPA) has significantly expanded its scope since its launch in 2005, incorporating immunohistochemistry data from normal and cancer tissues^7^. By 2015, HPA incorporated transcriptomic data of 32 normal tissue types, and had significantly expanded its scope to encompass 20,456 antibodies across 44 normal tissues^8^. The latest version includes data from over 27,000 antibodies, targeting a wide range of proteins.

However, despite the advantages of antibody-based methods in providing localized protein information, only proteins with highly specific antibodies could be reliably characterized, leaving out the majority of the human proteoforms understudied^9,10^. Mass spectrometry (MS) has emerged as an alternative tool for generating a comprehensive proteomic map of human tissues due to its ability to simultaneously analyze multiple proteins and its unbiased detection capabilities compared to antibody-based assays^11^. In 2014, two mass spectrometry (MS)-based human proteome drafts claimed the identification of approximately 85% of proteins encoded by human genes across 30 tissue types and cell lines^12,13^. Wang et al. performed a label-free MS analysis on 29 tissues^14^ and systematically analyzed the protein-per-mRNA by the combination of protein and previously generated mRNA data^8,15^. Jiang et al. recently quantified 12,027 proteins across 32 tissue types using Tandem Mass Tag (TMT)-based quantification^16^. While these studies have significantly advanced tissue protein identification, they primarily focus on 30 major tissue types, leaving the proteomes of many other tissues uncharted. Additionally, the lack of comparisons between benign and cancerous tissues limits their applicability to pathological conditions.

Cancer can originate from various tissue types, and most therapies are administered systemically; therefore, identifying specific molecular changes in each cancer type is crucial for accurate diagnosis and the identification of anti-cancer drug targets. The Cancer Genome Atlas (TCGA) and the International Cancer Genome Consortium (ICGC) have generated extensive genomic and transcriptomic data^17,18^. In recent years, the Clinical Proteomic Tumor Analysis Consortium (CPTAC) reported proteomic datasets for multiple cancers, and integrated multi-omics data analysis^19,20^. Although these and many other studies in the literature have adopted MS-based proteomics technologies, which are continuously improving, technical challenges remain in harmonizing identifiers, ensuring data integrity, and mitigating batch effects^21^. In addition, there is a lack of comparison to normal tissues in multi-omics analyses.

In-depth profiling of the human body proteome across baseline and tumor tissues necessitates a high-throughput, sensitive, and robust MS approach. Data-independent acquisition (DIA) MS method has emerged as a more powerful strategy for large-scale quantitative proteomics profiling^22-24^. Here, we present a comprehensive pan-cancer and healthy human tissue proteome landscape using DIA-MS. This resource encompasses nearly all solid human tissues, body fluids, and major cancer types (**Figure 1**). We report a detailed spatial distribution of over 10,000 proteins across the human body in both healthy and cancerous states. Additionally, we identified numerous previously unreported missing proteins, undetected by both MS and affinity-based methods. This resource also uncovered multiple potential drug targets.

**Figure 1.**
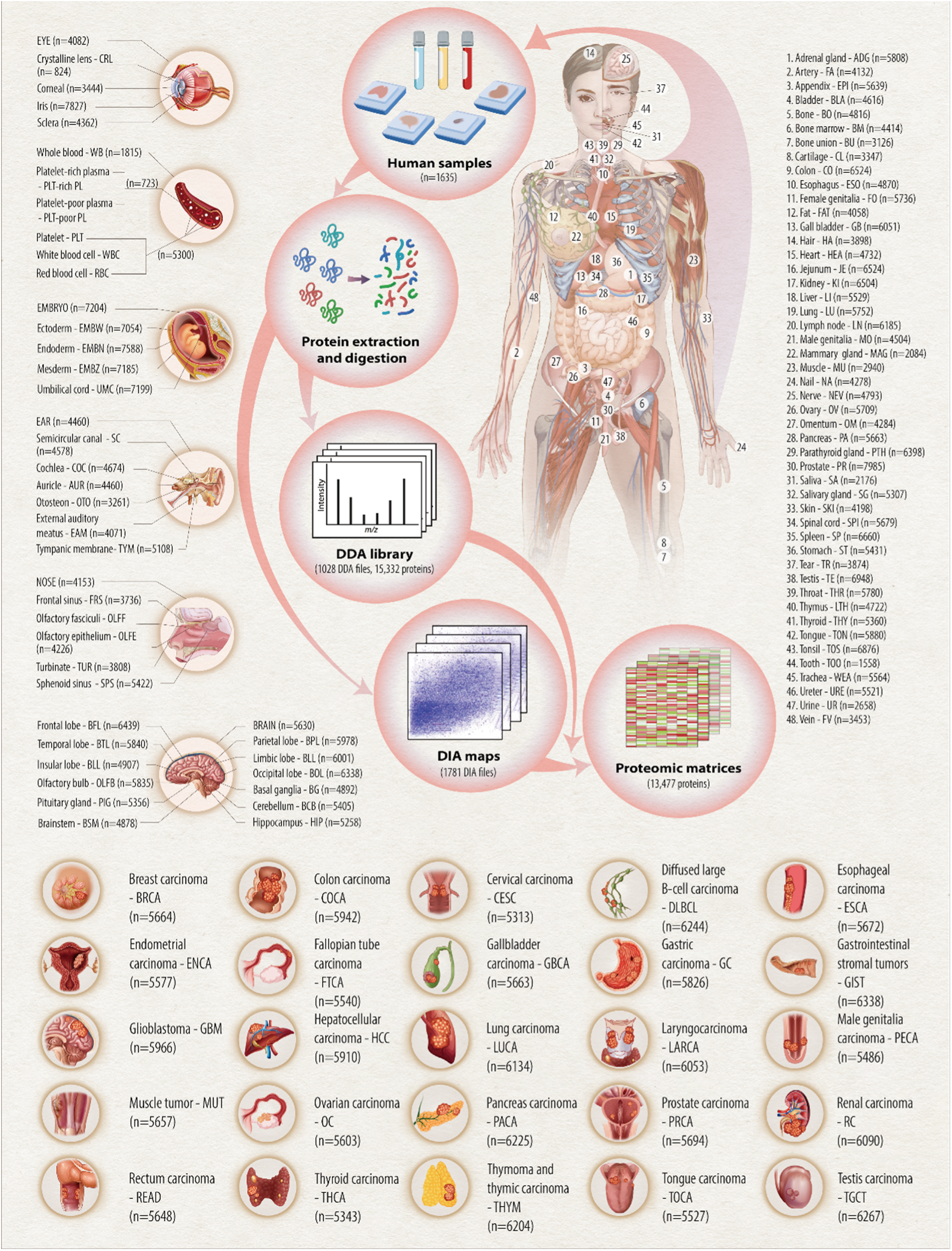
Overview of the human proteome draft. Figure 1 exhibits the types of samples contained in our data, with each sample information marked with a corresponding number. The specific sampling positions of the eye, blood, human embryo, ear, nose, and brain are listed on the left. 25 types of cancer are shown at the bottom. All samples are labeled with the full and abbreviated names of the organs/tissues/cancers, and the numbers in parentheses indicate the number of proteins in the database.

## RESULTS AND DISCUSSION

### Samples and proteomic profiling

To generate a comprehensive proteomic profile of healthy and cancerous human tissues, we collected 1635 samples from nine post-mortem adult donors, eight healthy participants, nine post-mortem fetal donors, and 519 cancer patients (**Table S1**). We first built a comprehensive spectral library using these samples (**Supplementary information (SI), Figure S1A**). A total of 15,322 proteins were identified in the spectral library, while 13,477 proteins were quantified across 1781 MS raw files (**Figure 1**). The proteomes displayed substantial heterogeneity across various tissue and sample types (**SI, Figures S2A, S2B, S2F, S2G**), with a high degree of reproducibility among replicates, underscoring the data quality (**SI, Figures S2C, S2D, S2E**).

T-distributed Stochastic Neighbor Embedding (t-SNE) analysis on the 1781 samples exhibited an orderly arrangement of fetal samples (F), tumor samples (T), paired non-tumor samples (NT), and normal samples (N) along the opposite direction of the x-axis (t-SNE1, **Figure 2A**), mirroring the degree of tissue differentiation. To elucidate the specific proteins and biological processes underpinning this transition, we conducted Wilcoxon rank-sum tests to identify the differentially expressed proteins across the F-T, T-NT, and NT-N groups. A total of 222 proteins were significantly differentially expressed across these three groups, including 187 downregulated proteins and 35 upregulated proteins, indicating either increased or decreased expression levels along the F-T-NT-N trend. Subsequent functional enrichment analysis of these proteins revealed that the upregulated proteins were predominantly enriched in RNA splicing and transport, as well as histone deacetylation (**Figure 2B**), suggesting a shared demand for rapid RNA transcription and splicing in both cancer and early embryonic development. RNA alternative splicing, a conserved process widely implicated in organ development, has been reported to be reactivated in various cancer types, influencing prognosis and growth rates^25^. Conversely, the downregulated proteins were primarily enriched in immune functions, particularly humoral immune response (**Figure 2B**). We propose that this downregulation may be associated with suppressed or incomplete humoral immunity in prenatal or tumor samples^26,27^.

**Figure 2.**
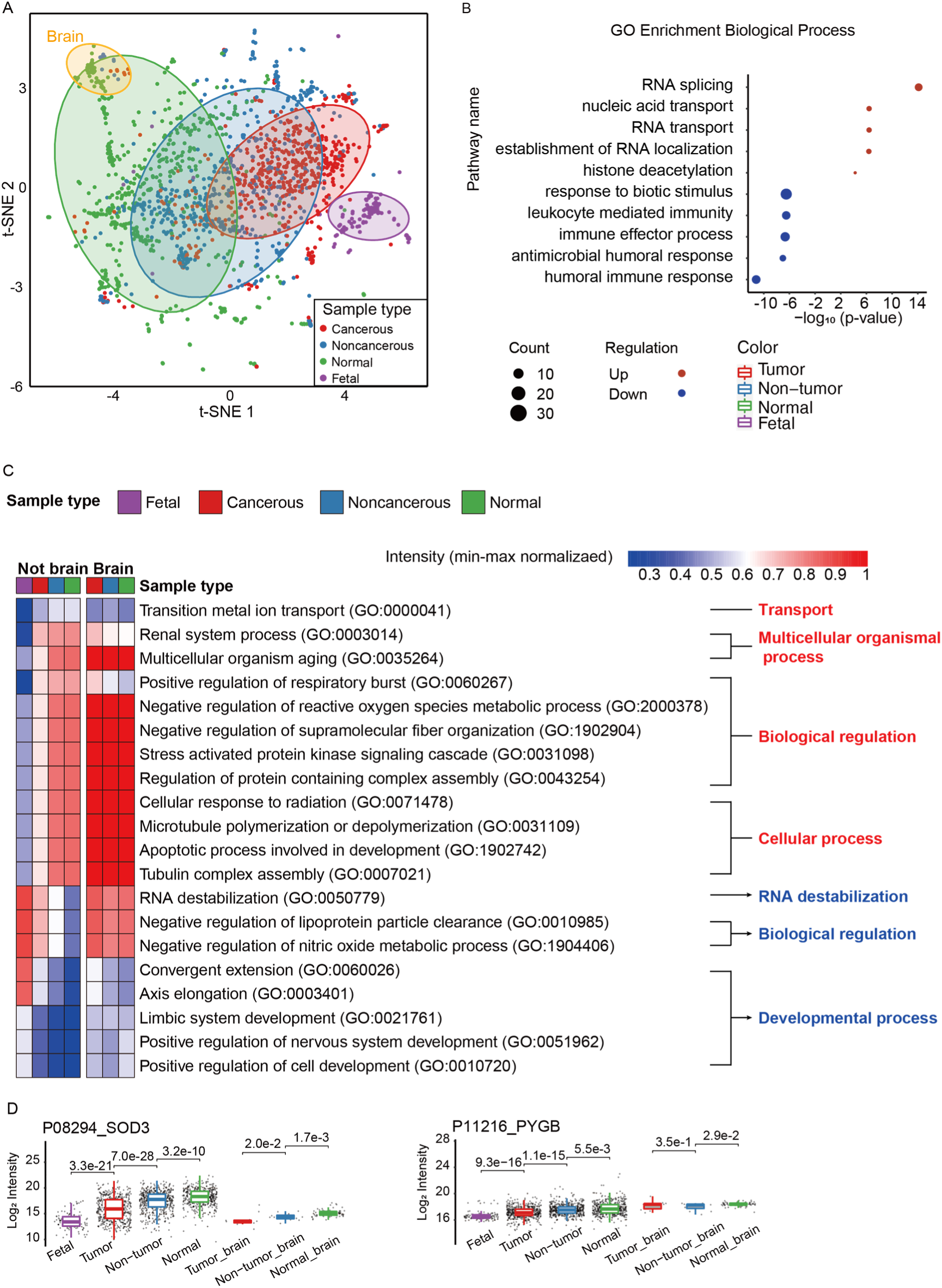
Proteome transformation in tissue development and oncogenesis. **A**, t-SNE plots of carcinoma, adjacent, normal and fetal samples. **B**, Functional enrichment for 222 proteins. **C**, GOBP gene set enrichment analysis for protein levels in different sample types of brain and non-brain samples. **D**, box plots exhibited 2 proteins of different expression patterns across sample types in brain and non-brain samples.

We observed that certain brain tumors and non-tumor tissues deviated from the F-T-NT-N trend, instead clustering together with their respective normal tissues. The uniqueness of gene expression evolution during brain organ development has also been reported. A comparative analysis of transcriptomes of seven organs, including the brain, across various species and developmental stages, revealed that gene expression during human brain development was more functionally constrained and exhibited the most pronounced differences in gene expression trajectories between species^28^. We compared GO abundances between brain and other tissue types across multiple sample types and found suppressed metabolism of reactive oxygen species and respiratory burst in the brain (**Figure 2C**). Superoxide dismutase 3 (SOD3), an antioxidant enzyme that catalyzes the conversion of superoxide radicals into hydrogen peroxide and oxygen and protects brain, exhibited a relatively lower level among human tissues, which is consistent with HPA proteome and transcriptome results^8^. However, metabolism-related pathways exhibited markedly higher activities in all brain sample types compared to other tissues, likely due to the brain’s substantial energy demands. Glycogen phosphorylase B (PYGB) is predominantly found in the brain as a rate-limiting enzyme in glycogenolysis. We found that PYGB consistently showed high expression levels in the brain (**Figure 2D**), which might be explained by its essential role in brain metabolism.

### Tissue specificity of protein expression across the human body

Through evaluation of the Euclidean distances of samples within major and sub-categories of tissues, along with the correlation within and outside category, we were able to evaluate whether the classification of tissue type is reasonable on the proteomic level. As shown in Table S3, all the tissue types showed significantly higher correlation within compared to the outside category, except for vagina, eye, cartilage, fallopian tube and pancreas, with the highest distance. We have further refined the category accordingly (**Table S3**).

To clarify the proteomic characteristics of different human tissues, we analyzed the relative protein intensities among roughly partitioned tissue types of normal samples by post-mortem anatomy, except for the body fluids and embryo samples due to the huge differences in proteomes (**Figure S3F and S3G**). A total of 497 samples representing 51 rough tissue types from nine patients were selected. Firstly, we visualized the similarity of proteome among tissue types. From the hierarchical clustering (h-clustering) tree (**Figure 3A**), we found that physiologically related samples, like cerebral cortex and spinal cord, were more likely to cluster together. A few exceptions include: 1) peripheral nerve (NEV) and fat; 2) uterus and ureter; 3) throat and salivary gland. We found samples of these tissue types are dispersedly distributed in the t-SNE map (**Figure S2G**), and the similarity of one sample would cause the similarity of the whole tissue type on one-dimension h-clustering, as most tissue types have few samples.

**Figure 3.**
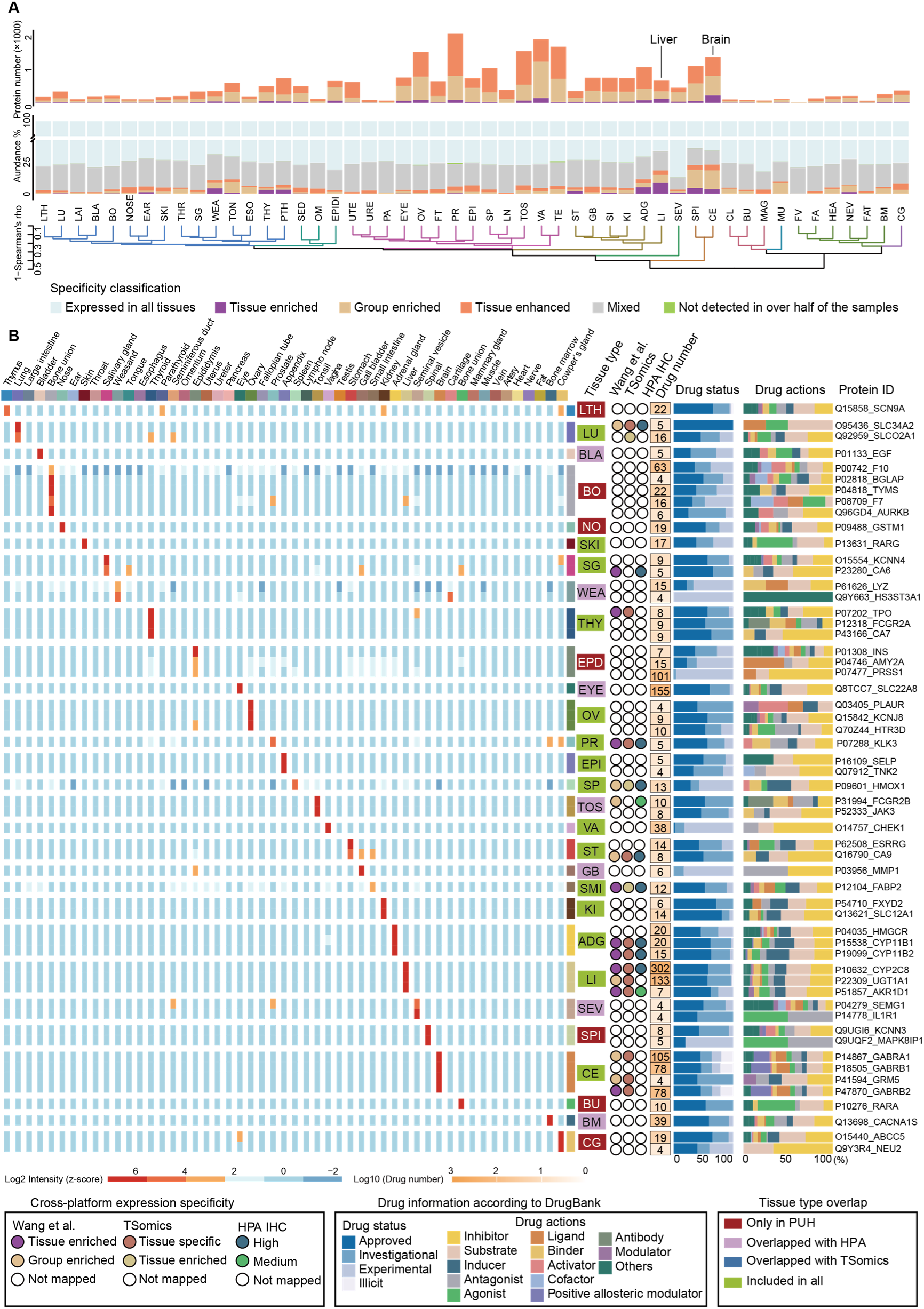
Tissue specificity of protein and drug targets. **A**, bar plot showing the protein numbers and abundance ratio of each specificity category in each tissue type. The bottom tree plot was generated by h-clustering the median abundance of each tissue type. **B**, the heatmap shows tissue specific drug targets. Only the top 5 highest enriched proteins are shown for each kind of anatomical classification. Tissue specificities of the targets are compared to previous human proteome drafts. For each drug target, the corresponding drug number and information are shown in colored bars.

To identify the enriched proteins for each tissue, we classified 12,392 proteins in 51 tissue types into six groups according to the HPA criteria^8^: not detected, tissue enriched, group enriched, expressed in all, tissue enhanced, and mixed, with the composition of these categories shown in Figure S4A and Table S3. The proteins expressed in all tissues constituted around a quarter of the quantified protein IDs, and three-quarters of the total abundance (**Figure 3A**). The functional enrichment of proteins specific to major tissue types is shown in Figure S4A and aligns with biological expectations (**Table S5**). As expected, proteins related to metabolism, synaptic function, gonadal and germ cell development are notably enriched in liver, brain, ovary and testis. Additionally, cilium-related pathways were enriched in both the seminal vesicle and vas deferens. We also identified common enrichment patterns across several distinct tissue types. For example, proteins involved in oxidation-reduction processes are highly enriched in metabolically active tissues such as the heart, muscle, brain, liver, kidney, and stomach (**Table S4)**.

Brain contained the highest number of tissue-enriched proteins, whereas the summed abundance ratio of tissue enriched proteins to all identified proteins was highest in liver (**Figure 3A**). The former has also been reported in previous studies due to the distinct biological function of brain^8^, whereas the latter has been ignored. As shown in Figures S4B and S4C, liver-enriched proteins ranked in the top 20% of the liver proteome, while brain-enriched proteins mostly ranked in the bottom 80% of the brain proteome. We inferred that the high abundance ratio of tissue enriched proteins might be related to the higher burden of metabolism as indicated by the dominant metabolic pathways enriched by liver-enriched proteins (**Figure S4D**). Most of the brain tissues were enriched in neural-related pathways (**Figure S4E**), suggesting that the enriched proteins could reflect the primary biological functions of the tissues.

Collectively, we have developed a more comprehensive view of the proteomic landscape across human tissues, highlighting their tissue-specific expression of proteins. These findings could have important implications for understanding the molecular mechanisms underlying tissue-specific functions and diseases.

### Tissue specificity of drug target proteins for understanding systemic drug response

Most drug targets are proteins^29^. A comprehensive map of the expression of drug-targetable proteins in various tissues and organs was suggested as a reference for the potential selection of drugs to avoid interaction with off-targets and to eliminate unintended systemic side effects^30^. We found 329 tissue enriched proteins were drug targets in DrugBank^31^, corresponding to 2536 drug molecules (**Table S4**). Compared to previous studies, seven tissue types of at least one tissue enriched drug target were found exclusively in our data, including the thymus, bone, nose, seminiferous duct, spinal cord, bone union, and Cowper’s gland (**Figure 3B**). We found a protein enriched in bone named osteocalcin (BGLAP), which was classified into tissue enhanced proteins in choroid plexus and intestine in HPA (**Figure 3B**). BGALP, a 10.96 kDa protein of 100 amino acids, is mainly produced by osteoblasts^32^. The misclassification by HPA likely stems from the lack of bone-derived samples in their dataset, where BGALP shows low transcription across tissues (<10 transcripts per million), with relatively higher levels in the choroid plexus and intestine. Beyond its structural role, bone is involved in energy metabolism and endocrine regulation. Initially linked only to skeletal development, osteocalcin is now known to affect systemic metabolism, reproduction, and cognition^32^. Our database offers a resource for further exploration of bone’s multifaceted biological roles.

To clarify how this resource enhances our existing biological or clinical insights, we searched for drug target proteins in the tissue enriched proteins and compared them with previous studies^8,14,16^. Liver had the most tissue enriched drug targets (**Table S4**), which might partly explain the high tendency towards liver damage in drug side effects. Cytochrome P450 2C8 (CYP2C8) is found to be highly enriched in liver tissue (**Figure 3B**) both in our data and some external protein expression datasets (HPA^8^ and GTEx^16^). This enzyme serves as the target for 302 drugs listed in DrugBank^31^, the majority of which are approved drugs on DrugBank spanning various therapeutic classes including antivirals (e.g., amodiaquine), metabolism-modifying agents (e.g., cerivastatin), and anticancer drugs (e.g., imatinib). In most cases, these drugs don’t directly interact with CYP2C8 to realize their pharmacological mechanism, but acting as inhibitors to substrates instead (**Figure 3B**), potentially causing liver toxicity, given the involvement of hepatic CYPs in the pathogenesis of liver diseases^33^.

Besides, we investigated possible organs impacted by off-target and validated through description in case reports. We interrogated the literately reported cases of side effect of the drugs targeting the top five enriched drug targets (**Figure 3B**). Specifically, we focused on tissues showed no direct drug response, excluding those with blood barriers or hormone effects (**Table S4**). We screened and found a clinical case of salivary gland side effects caused by Ritodrine aligning with our hypothesis. Ritodrine is a beta-adrenergic agonist which was historically used to treat uterine contraction and premature labor^34^. It activates Ca^2+^-activated K^+^ channels^35^ and inhibits plasma membrane Ca^2+^-ATPase^36^. The protein KCNN4, enriched in the salivary gland, forms the calcium-activated potassium channel involved in saliva secretion. Activation of KCNN4 by ritodrine likely leads to increased salivary gland secretion, consistent with reported cases of painless bilateral swelling of parotid and submandibular glands in some pregnant women receiving ritodrine hydrochloride treatment^37-39^. By looking back into this case, we demonstrated the potential utility of this database in conjunction with DrugBank for predicting systemic side effects. By interpreting reported clinical cases, we provided evidence supporting the value of this approach in understanding drug-induced adverse reactions.

### Comparative analysis between paired tumor and non-tumor tissue identifies key protein changes for oncogenic pathways

Due to the significant variances among sample types, tumor and their paired NT samples were subset to produce a pan-cancer proteomic landscape (**Figure S3A**). This is the first pan-cancer proteomic dataset processed and analyzed using the same pipeline and analysis platform including paired carcinoma and adjacent samples. We employed batch design to minimize the batch effect caused by multi-batches and multi-platforms^40^ and provided new aspects for the treatment of cancers. Quality control analysis exhibited high reproducibility and minimal batch effect after correction (**SI, Figure S3B-F**). We observed a great distinction of proteome between different types of carcinomas as shown by the t-SNE map (**Figure S3G**), especially for glioblastoma (GBM), diffuse large B-cell lymphoma (DLBCL), and hepatocellular carcinoma (HCC). Among the less discriminative cancer types, we found that gynecological carcinomas including breast carcinoma (BRCA), endometrial carcinoma (ENCA), cervical carcinoma (CESC), ovarian cancer (OC) and fallopian tube carcinoma (FTCA) were clustered together, while the cluster of archenteric carcinomas including colon carcinoma (COCA), rectum carcinoma (RECA), gastric carcinoma (GC) and pancreatic carcinoma (PACA) was distinguishable from the gynecological carcinoma cluster (**Figure S3G**). Considering the different germ layers of organ origin in these two clusters, we speculated that the distinct proteomic structures might correlate with patterns shared in early embryo development and oncogenesis.

In contrast to other large-scale tumor characterization studies, all of the tumor types included paired tumor (T) and non-tumor adjacent tissue (NT) samples, allowing us to explore divergent expression patterns between T and paired NT samples. We found that reproductive system neoplasms, including ENCA, BRCA, testis carcinoma (TGCT) and CESC, had the highest numbers of significantly differentially expressed proteins (DEPs), while digestive system cancers such as gall bladder carcinoma (GBCA), COCA, GC and HCC had low numbers of DEPs (**Figure 4A, Table S5**). GBM had the lowest number of DEPs, which might be associated with stricter functional constraints in brain. Cross-comparison of the divergent expression between T and NT in our data, TCGA and CPTAC datasets showed considerable consistency of the matched genes, and over 3000 proteins were coherently clarified as significantly upregulated or downregulated proteins in the three datasets (**SI**).

**Figure 4.**
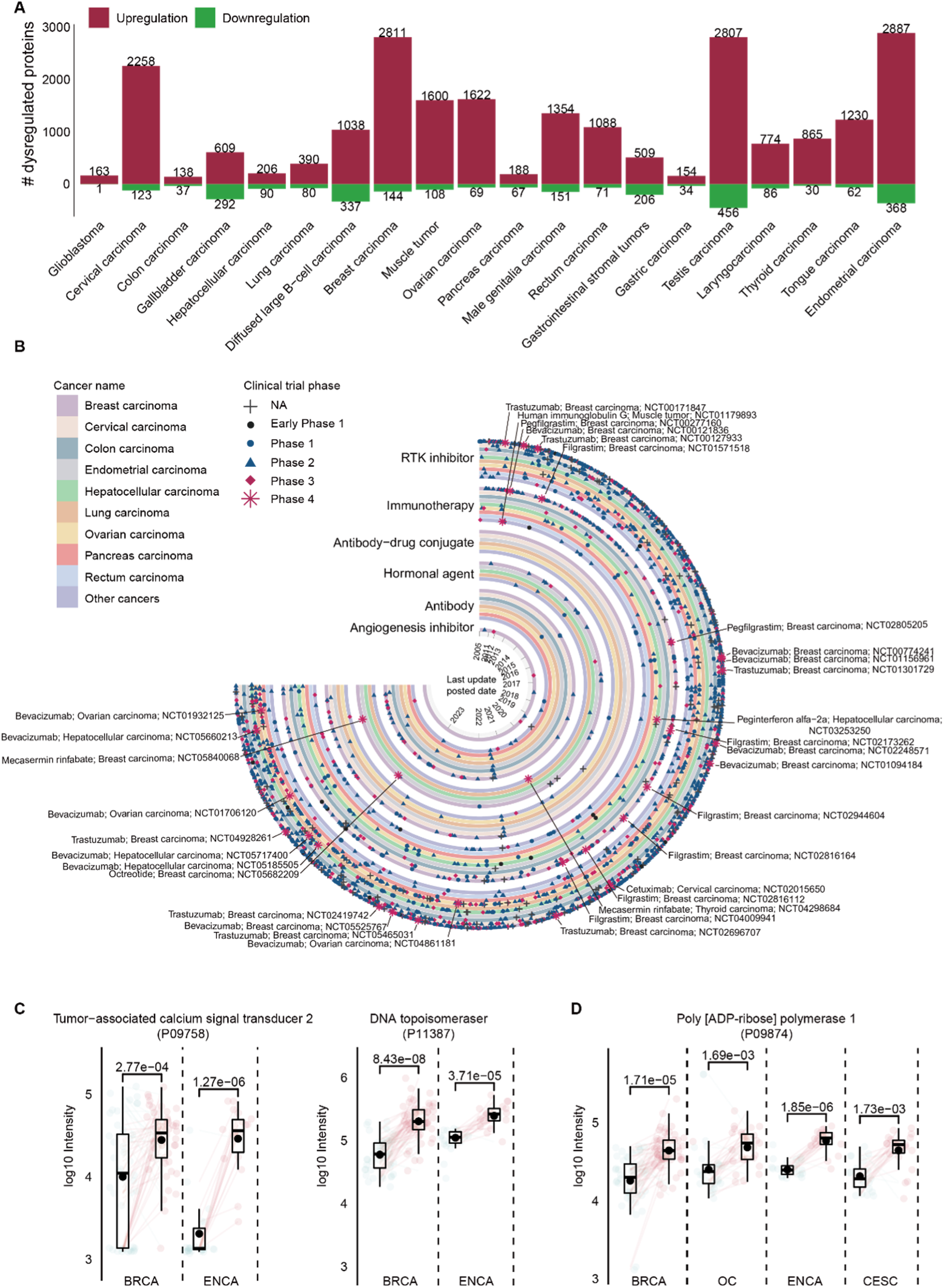
Pan-cancer analysis of dysregulated proteins between paired tumor and non-tumor samples. **A,** Carcinoma dysregulated protein. Red bars mark upregulations and green bars mark downregulations. **B**, DEPs from all cancer types mapped to targets and clinical trials. The clockwise direction represents the start time of clinical trials, circular layers indicate drug types, track colors correspond to cancer types, and symbols denote trial phases. Scatter plots show the expressions of the target of sacituzumab govitecan (**C**) and olaparib (**D**). Points in red and green represent carcinoma and adjacent samples respectively. Box plot shows protein expression; center lines indicate the medians and center points indicate the means; box limits represent the first and third quartiles; whiskers extend 1.5 times the interquartile range from the quartiles to outliers; translucent scatters show all sample points. Colored lines connect the carcinoma-adjacent pairs, while red lines mean up-regulations and green lines mean down-regulations. BH-adjusted *P* values (two-tailed paired t-test) are indicated by stars, * p<0.05, ** p<0.01, *** p<0.001.

In most cases, upregulated DEPs were unique to a single cancer (cancer-specific) or shared between two cancers (**Figure S7A**). Among the 25 cancer types, TGCT and DLBCL shared the most upregulated DEPs (n =202, **Figure S7A**), followed by BRCA and ENCA (n = 122, **Figure S7A**). A total of 56 DEPs were upregulated in at least 12 cancer types (**Figure S7B**), including some well-known oncogenes like Signal transducer and activator of transcription 1 (STAT1) and DNA topoisomerase 2-alpha (TOP2A) (**Figure S7C**). Consistent with CPTAC pan-cancer cohort^41^, upregulation of PLOD2 was observed at the pan-cancer level (**Figure S7C**), indicating its potential as a pan-cancer prognostic biomarker. The generalizability of these aberrant expression patterns suggests their importance in oncogenesis.

We found some of these cancer-specific DEPs have been reported associated with both tissue differentiation and oncogenesis (**SI**). To validate these specific protein dysregulations, we performed parallel reaction monitoring (PRM) analysis for 14 of these cancer specific proteins on all carcinoma samples and most normal samples, among which five were consistent with the DIA quantification result. ADAMDEC1 (ADAM-like decysin-1) was found to be specifically upregulated in DLBCL both in our DIA and PRM data (**Figure S6C**). Consistently, Zhu et al. found that ADAMDEC1 was upregulated in DLBCL, and its expression levels correlated with tumor progression and poor patient survival^42^.

### Pan-cancer dataset facilitates drug repurposing and target discovery

To further explore the potential therapeutic use of the DEPs, we mapped the 7641 DEPs found in all cancer types to the protein targets of biotechnical and large molecule drugs in DrugBank^31^ and ongoing clinical trials. A total of 236 DEPs were identified as drug targets corresponding to 289 drug candidates, with 2,089 clinical trials linked to 40 drugs that target a mere 81 DEPs (**Figure 4B, Table S6, SI**). These findings underscore the vast potential of this dataset for the discovery of novel drug targets. Here are three application scenarios.

The first scenario is the targeted therapy of cancer. The dysregulations of certain protein targets suggested viable and approved therapy strategies for cancers. Usually, there are some well-studied or approved drugs for certain protein targets in cancer treatment, but their efficacy in other cancer types remains understudied. Actually, it is rational that a verified upregulated protein across different cancer types might also be a potential drug target for the approved drug, even if in other types of cancers. For example, Trodelvy (sacituzumab govitecan) is an antibody-drug conjugate (ADC) agent that combines humanized anti-trophoblast cell-surface antigen 2 (TROP-2) antibody with the DNA topoisomerase 1 (TOP1) inhibitor SN-38. It was approved by the FDA for the treatment of triple-negative breast cancer (TNBC) by inducing DNA damage-mediated cell death. In our dataset, we observed upregulation of both the antibody target TROP-2 and the drug target TOP1 in ENCA and BRCA (**Figure 4C**). Compared to BRCA, few drugs were approved or under study for ENCA on the ClinicalTrials.gov database (by June 2, 2023). This suggested the potential of Trodelvy as an alternative drug for ENCA treatment, which is supported by a phase I/II clinical trial (NCT01631552)^43^. Another example is Poly [ADP-ribose] polymerase (PARP) inhibitor. It is well known that PARP1 was upregulated in BRCA and gynecologic cancers including OC, ENCA, and CESC (**Figure 4D**). Lynparza (olaparib) is a PARP inhibitor particularly selective for PARP1 and PARP2. Olaparib has been approved for the treatment of OC, FTCA, and BRCA. Although it is not currently used for the treatment of ENCA, associated phase I and II clinical trials have shown promising responses^44,45^.

The second is regarding receptor tyrosine kinases (RTKs). We evaluated ten oncogenic signaling pathways presented by Sanchez-Vega et al. in 2018^46^, with frequent and key genetic alterations reported in previous TCGA publications, and focused on pathway members likely to be drug targets. In our matched protein-gene pairs, the RTK-RAS pathway has the highest number of DEPs (**Figure 5A, Table S8**), and the highest median frequency of alterations (46% of samples) across all cancer types (**Figure 5A**)^46^. Given the potential for co-dysregulation of the RTKs might develop into drug resistance in the context of a single RTK inhibitor, we explored the co-occurrence of RTK dysregulation in our data (**Figure S5D**). Among all the RTKs, Epidermal growth factor receptor (EGFR) is the most commonly dysregulated RTK and most frequently co-occurred with insulin-like growth factor 1 receptor (IGF1R), which is the second most frequent alteration. The combination inhibition of EGFR and IGF1R has been extensively studied in the context of cancer therapy in a variety of cancer types^47,48^. However, clinical trials of dual EGFR/IGF1R inhibition have had mixed results, one reason to explain this would be the cancer heterogeneity.

**Figure 5.**
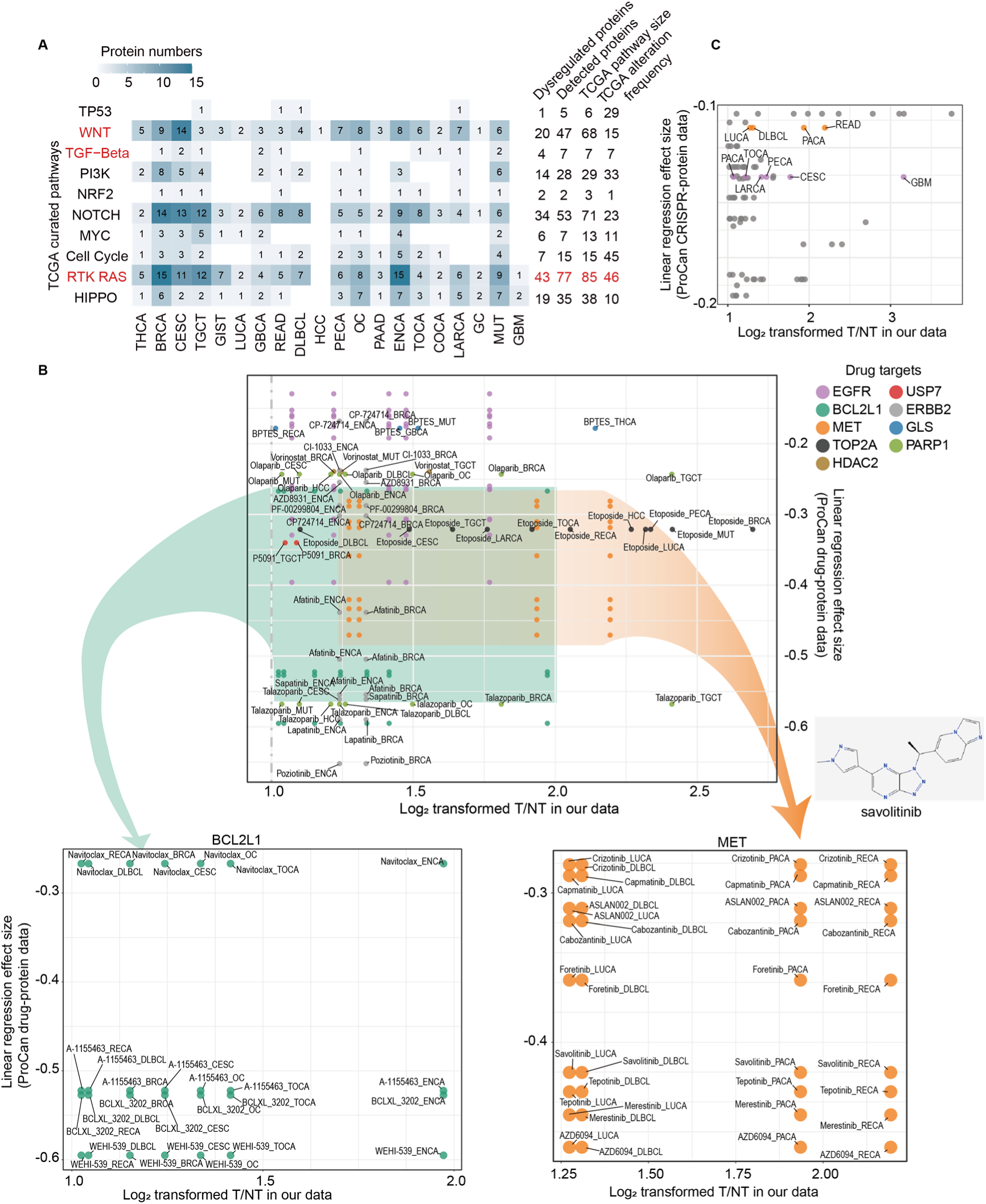
Potential drug targets and drug uses. **A,** Number of dysregulated proteins in TCGA curated pathways of each kind of cancer. Druggable carcinoma dysregulated targets mapped to ProCan-DepMapSanger drug-protein (**B**) and CRISPR-protein (**C**) association data. The color of a point represents its gene name. The drug-protein association results of BCL2L1 and MET are additionally shown.

To further investigate the correlation between dysregulations of the RTKs and drug response from a macro-view, we mapped our data to ProCan-DepMapSanger proteomic map^49^. By correlating the proteomic profiles of the cell lines with their drug response data and CRISPR-Cas9 gene essentiality data in 949 cell lines using linear regression, Gonçalves et al. were able to predict drug response and CRISPR-Cas9 gene essentiality based on the protein intensities^49^. Then we selected RTK proteins significant in the prediction of both drug response (beta < -0.1, FDR < 0.01) and upregulation (T/N fold change ≥ 2, B-H adjusted p-value < 0.05) in corresponding cancers (**Figure 5B**). In these mapped combinations, a higher absolute value of beta suggested stronger sensitivity of cell lines in a cancer type to a drug when the protein was more abundant, which is more favorable in the case of targeting upregulated DEPs. We found high expression of MET and BCL2L1 have been related to the higher potency of Navitoclax, a small molecule inhibitor of Bcl-2 family proteins in colon and rectum carcinoma cell lines and pan-cancer cell lines, respectively^49^. BCL2L1 and MET were also upregulated in RECA in our data (**Figure 5B**), which suggested the viability of this drug in the treatment of RECA. Besides, over-expression of MET has been reported in a variety of tumor types including colon cancer, pancreatic cancer and lung cancer^50^, which is consistent with our results (**Figure S5D**). MET-targeting drugs like Savolitinib and Tepotinib also exhibited higher potency at higher MET expression levels in pan-cancer cell lines including RECA cell lines^49^. The correlation between MET and RECA was further proved by the regression of MET protein intensity and CRISPR-Cas9 gene essentiality in RECA cancer cell lines (**Figure 5C**). In summary, we presented the clinical application of the pan-cancer proteomic database by focusing on the curated RTK/RAS signaling pathway, the most frequently dysregulated and mutated oncogenic pathways in 25 cancer types. Through integration with drug response and CRISPR gene essentiality data, we proposed the possible therapeutic value of targeting MET in RECA.

The third is to minimize drug adverse effects. Using the proteome of normal tissues from the whole body as a global map, we could select drug targets locally enriched in the tissue of lesions, thus minimizing systemic side effects. We filtered locally enriched DEPs (LEDEPs) for each tissue type by overlapping its tissue enriched proteins in the normal samples and the DEPs of the corresponding cancer types. A total of 121 LEDEPs were found in eight tissue types (**Table S9**). Some of the LEDEPs are well-studied, like Wilms tumor protein 1 (WT1)^51^ and Napsin-A (NAPSA)^52^, while most are yet to be noticed and validated. Functional analysis for these proteins suggested functional abnormality and distinct oncogenic signaling pathways, providing potential targets with high tissue specificity and aberrant expression for ADC drugs (**Figure 6, SI**).

**Figure 6.**
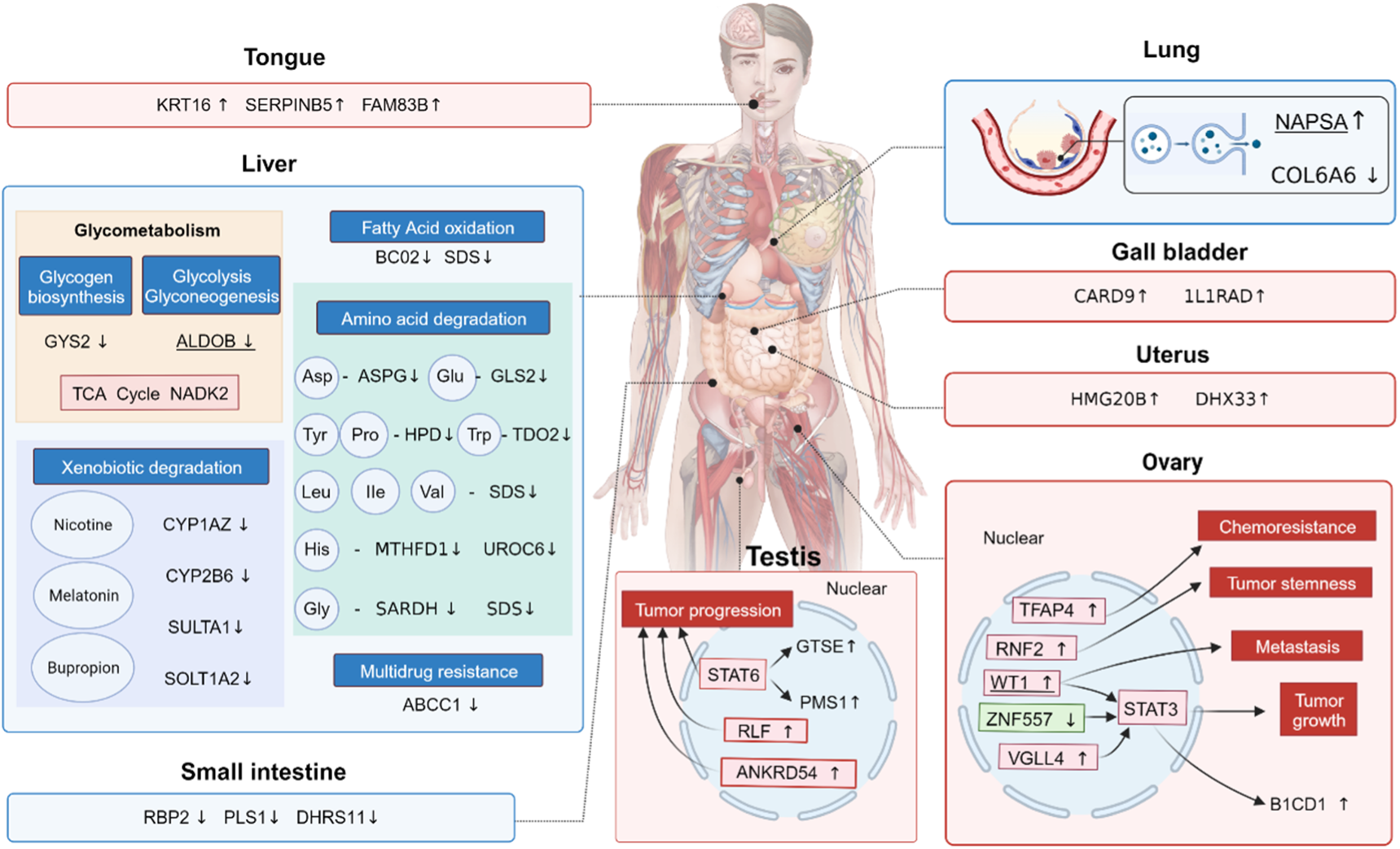
Sketch of locally enriched dysregulated proteins. A hypothetical systems view of the locally enriched DEPs (LEDEPs) in multiple organs. Up and down arrows stand for significantly up- and down-regulated in tumor compared to paired non-tumor samples. Blue and red blocks stand for activated or suppressed biological activities inferred by LEDEPs.

## CONCLUSIONS

This study described a comprehensive proteomic landscape for human tissues, containing normal autopsy samples from elderly individuals, body fluid samples, fetal samples, and carcinoma samples using the state-of-the-art high-resolution MS. This database extended the present knowledge about the protein evidence, completed as well as detailed the protein expression characteristics throughout the human body, and facilitated the discovery and evaluation of drug targets.

Our analysis of the normal body tissues portrayed the proteomic map for tissues that have merely been studied and not included in the existing databases of human gene expression like Cowper’s gland, ear and nose. Sub-organ tissues and the same tissue types at different anatomical positions were able to be further analyzed. Besides, as most drug targets are proteins, this comprehensive proteome draft enabled a fully systematic evaluation of the drug response. Our analysis for the carcinoma and paired adjacent samples filled the gap of lacking a considerable amount of pan-cancer data from the same analysis platform. We analyzed the commonalities and differences among the 25 types of carcinomas we collected. Shared DEPs between well-studied carcinomas like BRCA shed light on the medication of less-studied carcinomas like ENCA. Cancer-specific DEPs might greatly relate to the tissue specificity of the in-situ organs, as TGCT has the most cancer-specific DEPs. From the most frequently dysregulated and mutated oncogenic pathway RAS-RTK, we found possible drug usage in understudied cancer types and new drug combinations for refractory cancers. Through the combination of analysis of normal and neoplastic tissues throughout the human body, we were able to find locally enriched dysregulated proteins defined as LEDEPs. These proteins simultaneously possess the tissue traits and oncogenic alterations. Most LEDEPs are downregulated probably due to the dysfunction of the in-situ organs, while NAPSA, WT1 and PMS1 were upregulated in their respective organs. The former two are widely reported as the diagnostic and prognostic markers in lung and ovarian cancer, while we propose that they are also ideal as the drug targets with minimal systemic side effects.

This study also has some limitations. First, normal autopsies were mainly from a male and a female, as we have a very detailed partition of human tissues, limiting statistical power for sex- and age-related analyses. Besides, normal tissue types except for body fluids, hair and nail were only from elderly individuals, whose organs might have been affected by age-related degeneration. For example, normal mammary gland detected fewer proteins than the BRCA adjacent samples and reported numbers, probably because the mammary gland is degenerated and highly fibrosed in elderly women. Second, the majority of samples followed the two-step PCT protocol^53^, while we modified the protocol for bones, body fluids, cochlea, and olfactory epithelia samples due to the sample characteristics, which might cause manual interferences. Third, the sample size is not equal in the roughly partitioned tissue types. However, highly specialized tissues can be very robust according to the previous studies, while those mildly specialized would be affected. Despite all these limitations, this work presented a valuable resource for biochemical and biomedical research. We also have an obligation to have cell-type level quantitation data for these tissue types in the future.

## ACKNOWLEDGEMENTS

We would like to express our gratitude to the National Key R&D Program of China (Grant No. 2021YFA1301600, 2022YFF0608403). We are also grateful to the patients participated in this project for their donation of body or body tissue for scientific research. Additionally, we would like to thank the support teams of Bruker, FragPipe and DIA-NN for their assistance in acquiring and analyzing MS raw data, and we also appreciate the advanced computing services provided by Westlake University High-Performance Computing Center.

## AUTHOR CONTRIBUTIONS

T.G. and L.Y. conceived and designed the project. L.Y. and W.J. conducted the experiments, analyzed data, and wrote the manuscript. M.L., S.L., T.L. and N.X. assisted in sample preparation and MS data acquirement. Y.L., F.X., F.G., D.L. and N.F. conducted the autopsy, collected, and labeled the autopsy samples. T.L. and X.Z. procured the clinical samples and information. K.C. verified the pathological information of clinical samples. M.L. and J.A. assisted in the analysis of public database. W.G., Y.C., Z.F. and Z.N. assisted in building the data portal. F.Y., G.C.T. and A.N assisted in generating spectral library. B.C., M.S., and R.A. assisted in optimizing the data analysis workflow. Y.W., Y.Z. and R.A. helped with manuscript writing and editing. L.M. and L.L. proposed a list of tissue types for sample collection. T.G., Y. Z., Y.L. and T.L. supported and supervised the project. T.G. revised the manuscript. All the authors read and approved the manuscript.

## ETHICS

This research project has received approval from the relevant ethics committee or institution and has been conducted strictly in accordance with ethical guidelines. In this study, we respected and protected the rights and privacy of participants and ensured the confidentiality of their personal information. The autopsies were obtained from the Dalian Medical University (IRB: Dalian Medical 2019-05) and Shanghai Jiao Tong University (IRB: 2022A-01). The fetal autopsies came from the Southern Medical University Shenzhen Hospital-Gynecology (IRB: LLSCHY 2019-10-36). The paired tumor and non-tumor samples were obtained from the Harbin Medical University (IRB: KY2019-08, KY2023-03, KY2019-16).

## DATA ACCESSBILITY

All the qualitative and quantitative data utilized for analysis in this paper are openly accessible and fully available. Due to the limited publication length, the proteome data of some specific tissues or carcinomas were shown and discussed. To better visualize and present the entire database, we built a website (https://db.prottalks.com/), supporting both protein-centric and tissue-centric queries. Specifically, pairs of interested tissues can undergo differential expression analysis online, and the DEPs are shown in an instant heatmap. Raw data are also available for download at https://www.iprox.org/ (IPX0003578000).

## SUPPLEMENTARY TABLES AND FIGURES

**Supplementary Table 1. Sample origin information. A.** Information of non-carcinoma originated samples. The identifier, age, gender and disease of patients or volunteers, and the storage method of biological specimens have been listed. **B.** Information of cancer patient originated samples. The identifier, age, gender, tissue name, anatomical classification and corresponding tissue specific library name, and the origin center of patients have been listed. **C.** Statistical results of cancer patients. We are including patient number, gender ratio, mean and standard deviation of ages for each kind of cancers.

**Supplementary Table 2. DDA data identifications**

**A.** Identification of proteins in tissue-specific libraries. **B.** Identification of missing proteins in libraries. **C-D.** Statistics on missing proteins. **E.** List of missing proteins.

**Supplementary Table 3. Tissue specificity.**

**A.** Correlation, coefficient of variation (CV) and Euclidean distance within each of the major tissue types along with correlation between a specific tissue type and other tissue types. **B.** Median protein intensities in various tissue types. **C.** Classification of tissue specificity for each protein.

**Supplementary Table 4. Druggable tissue-enriched proteins information.**

**A.** Intensity matrix of tissue-enriched proteins (Z-score). **B.** Comparison of tissue-enriched proteins with external specificity databases. **C.** Search results of tissue-enriched proteins in DrugBank. **D.** Integrated research results by proteins. **E.** Statistics on drug status. **F.** Statistics on drug actions. **G.** Intensity matrix of the top 5 enriched proteins with at least three associated drugs. **H-J.** Subsets of **B**, **E**, and **F**.

**Supplementary Table 5. Differential expression analysis.**

**A.** Results of differential expression analysis between paired tumor and non-tumor samples for each cancer type. **B.** Differentially expressed proteins filtered by |fold-change| not less than 1.5 and adjusted p-value < 0.05.

**Supplementary Table 6. Cross-reference to external cancer datasets.**

**A.** Overlap of our data, TCGA, and CPTAC dysregulated proteins datasets. B. DEPs in HPA prognosis dataset. C. Overlap of **A** and **C**. D. Statistics of **B**.

**Supplementary Table 7. Cancer-specific proteins.**

**A.** Cancer-specific dysregulated proteins.

**Supplementary Table 8. Cancer DEPs associated pathways and drugs.**

**A.** DEPs detected in TCGA curated pathways. B. DEPs validated in ProCan-DepMapSanger proteomic map drug response dataset. C. DEPs validated in ProCan-DepMapSanger proteomic map CRISPR-Cas9 gene essentiality dataset.

**Supplementary Table 9. LEDEPs.**

**A.** Locally enriched dysregulated proteins with their enriched tissues, median expression intensity, and results of DEA.

**Figure S1.**
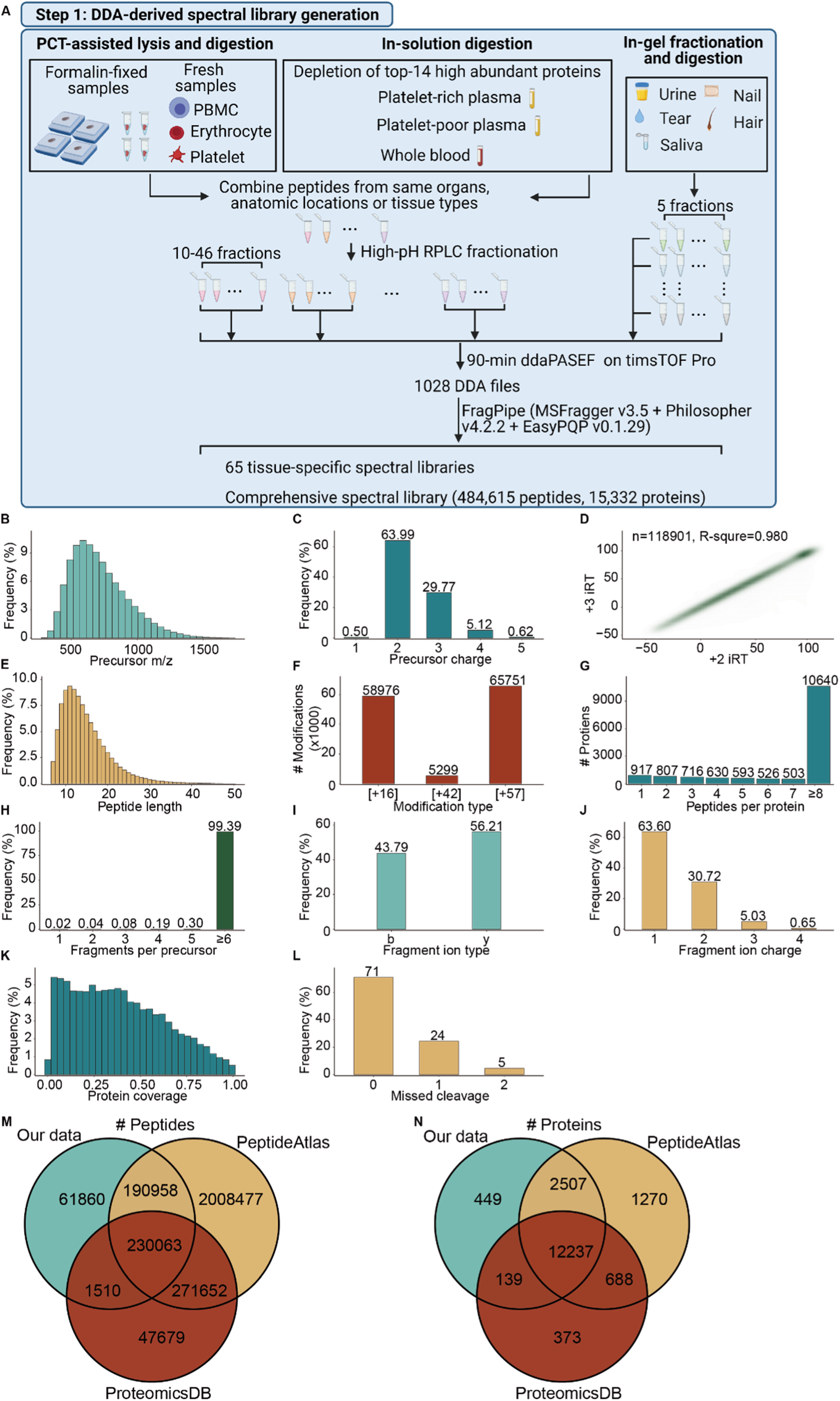
A view for quality of the spectral library. Percentage of precursor m/z (**A**), precursor charge (**B**). **C,** Pearson correlation of iRT values between +2 and +3 precursors with the same peptide sequence. **D,** Percentage of peptide length. Number of modifications (**E**), peptides per protein (**F**). Percentage of fragments per precursor (**G**), fragment ion (**H**), fragment ion charge (**I**), protein coverage (**J**) and missed cleavage ratio (**K)**. Overlapped peptides (**L**) and proteins (**M**) between our data, PeptideAtlas, and ProteomicsDB datasets.

**Figure S2.**
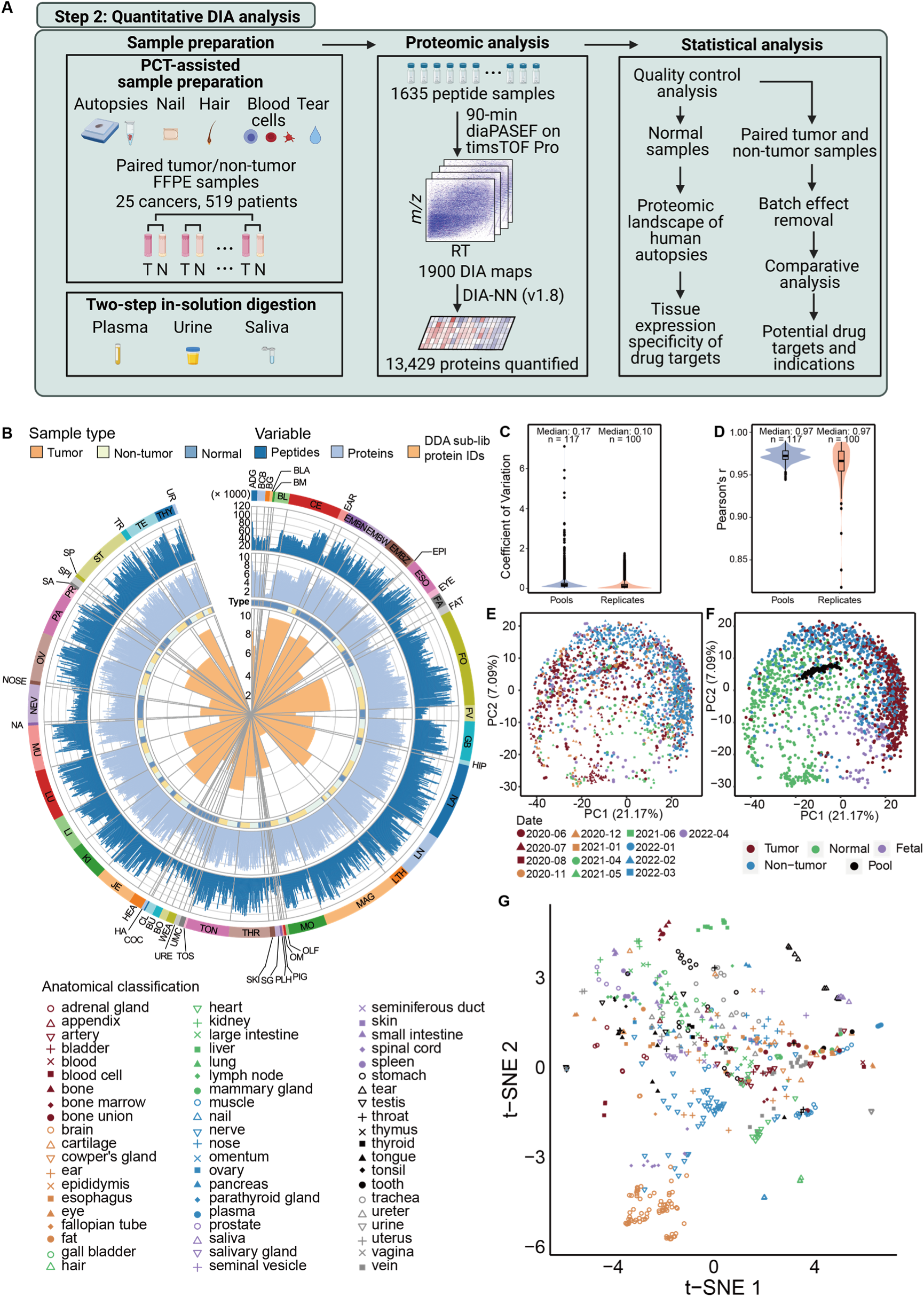
Overview of the analysis workflow and quality of the quantitative proteomic data. **A,** Workflow of DIA quantitative data analysis**. B**, Number of identified proteins in ddaPASEF data and quantified proteins in diaPASEF data. Each radius in the figure represents one sample. The radar plot in the center shows the number of proteins identified in the relative tissue-specific library for each sample. From the center of the circle, the 1^st^ circle shows the sample type, whether it is a tumor (T), tumor paired non-tumor sample (NT), or normal sample (N). The 2^nd^ circle represents the number of proteins identified in the sample, and the 3^rd^ circle represents the number of peptides identified in the sample. **C** and **D,** CV and Pearson’s correlation of pool and replicates. **E** and **F**, PCA plots showing the distribution of all samples, labeled with glass capillary (D) and sample types (E). **G**, t-SNE plot showing the distribution of normal samples, labeled with tissue types.

**Figure S3.**
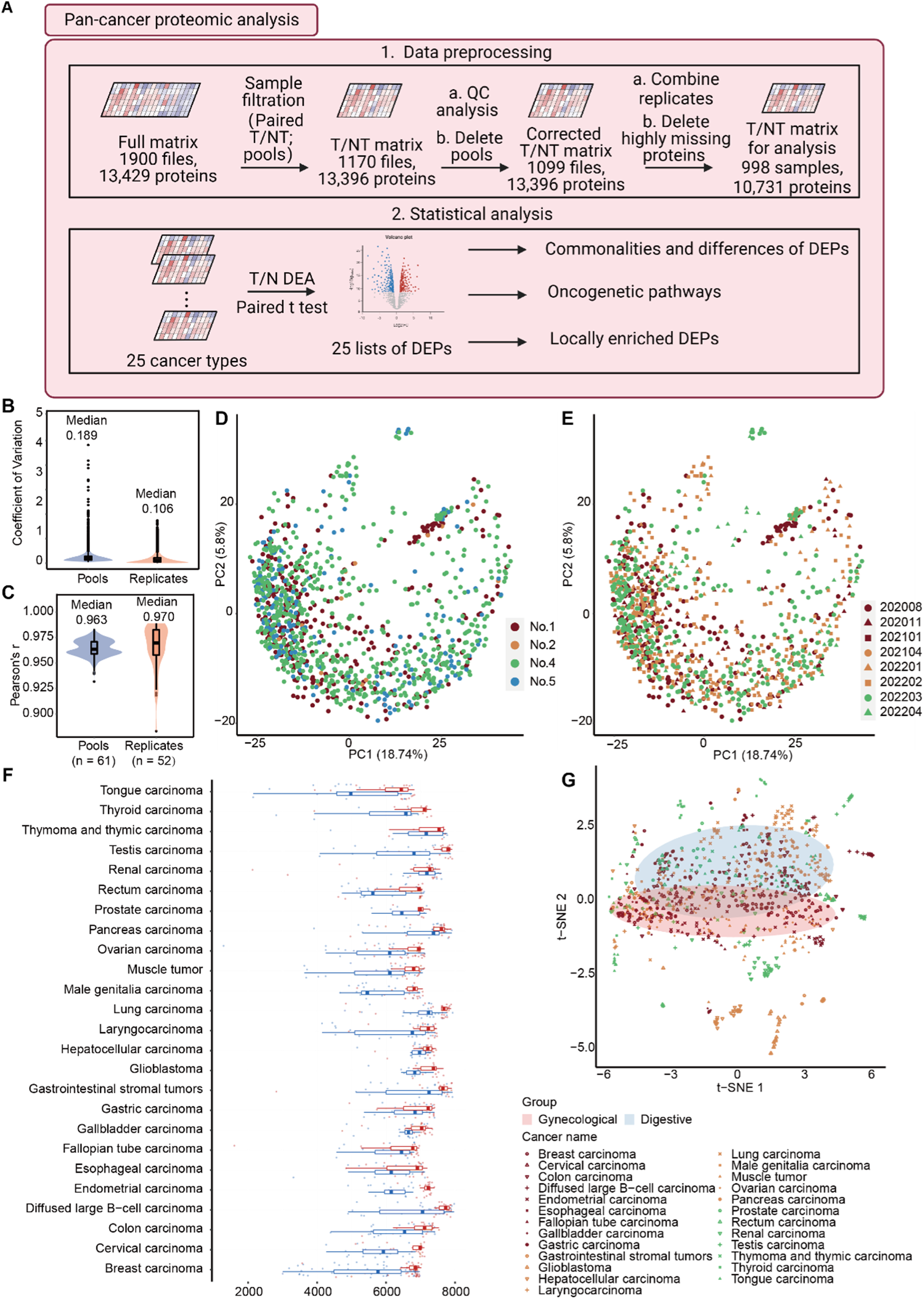
Overview of the pan-cancer quantitative proteomic data. **A,** Workflow of pan-cancer analysis**. B** and **C,** CV and Pearson’s correlation of pool and replicates. **D** and **E**, PCA plots showing the distribution of clinical samples, labeled with acquisition months (D) and glass capillary (E). **F**, Bar plots showing the numbers of identified proteins the tumor and non-tumor samples in 25 cancers. **G**, t-SNE plot showing the distribution of clinical samples, labeled with cancer types.

**Figure S4.**
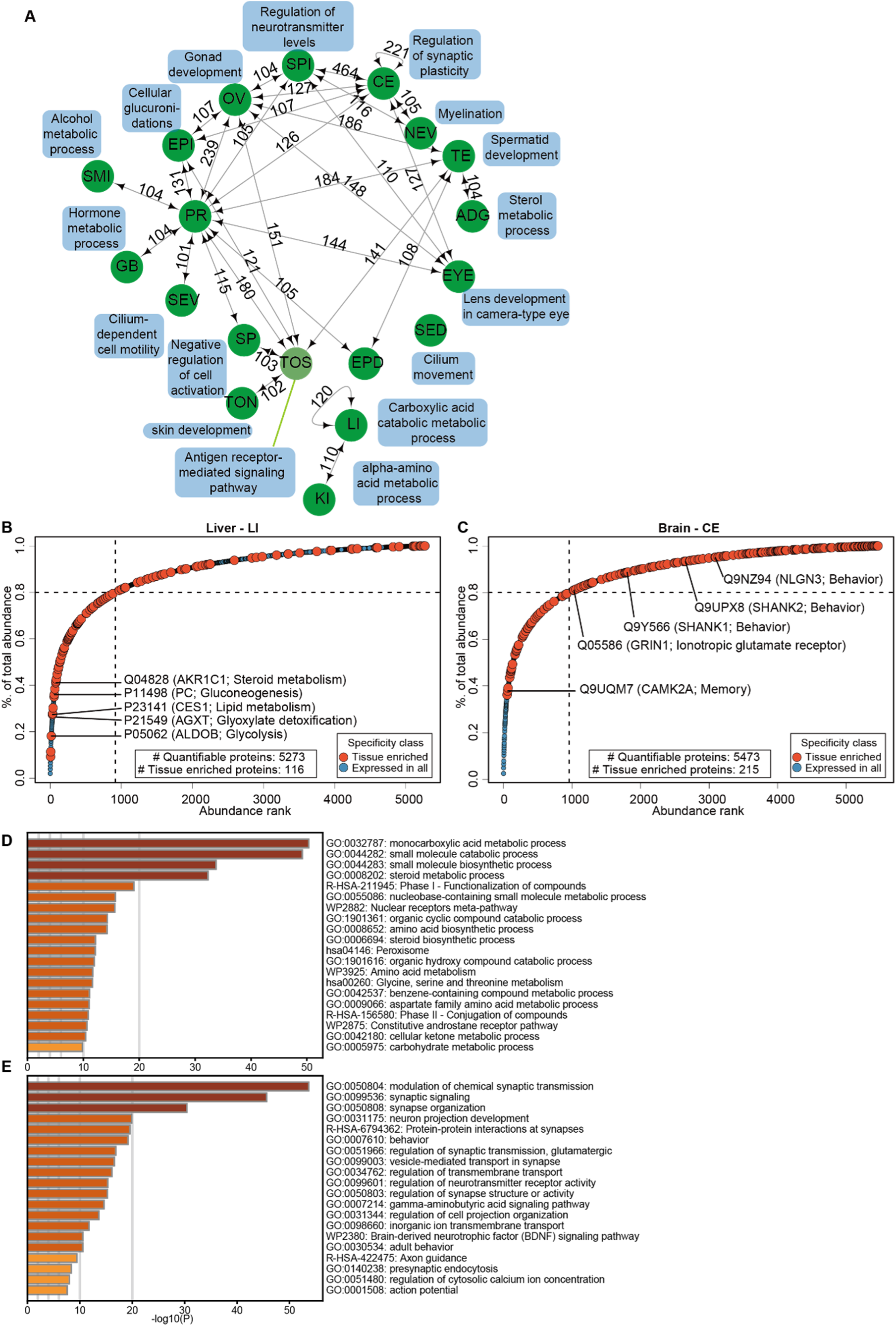
Tissue enriched protein and pathways. A, network showing the number of tissue enriched and group enriched proteins for each tissue type. Edge standing for two linked nodes (tissue types) shared more than 100 group enriched proteins. Mostly enriched GOBP term (by adjusted p-value and gene ratio) for each tissue is showing outside the network nearby the tissue node. B, C, dot plots showing the accumulative curves of protein abundance in liver (B) and brain (C). D, E, term enrichment for tissue enriched proteins in liver (D) and brain (E).

**Figure S5.**
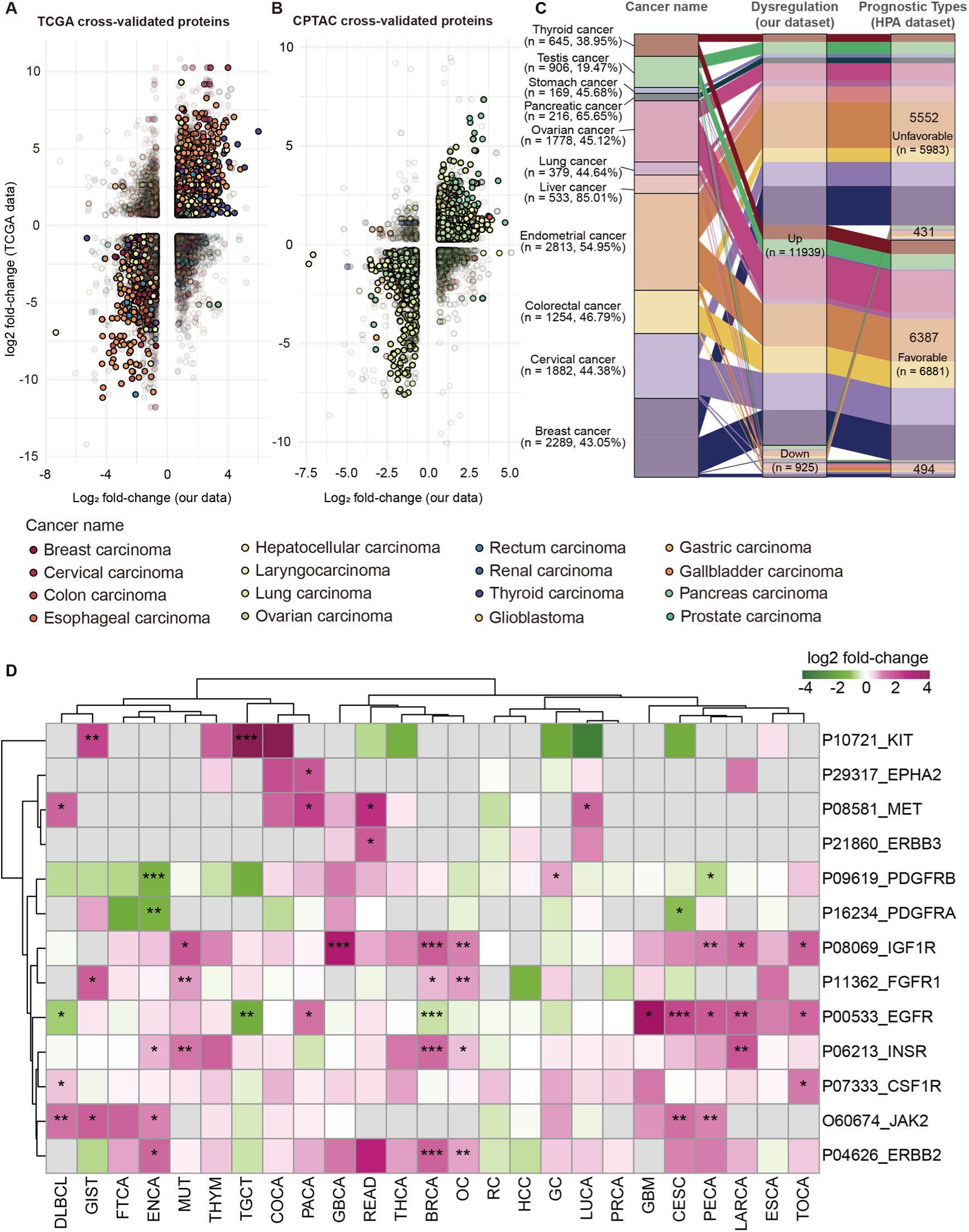
Dysregulated proteins in our data, TCGA and CPTAC. **A, B,** cross-validation in TCGA and CPTAC dataset. **C,** Overlap with HPA favorability dataset. **D**, RTKs dysregulations. The color of a square represents fold-change (log2 transformed) and non-detected cases are colored in grey. The BH-adjusted P values are indicated by stars, * p<0.05, ** p<0.01, *** p<0.001.

**Figure S6.**
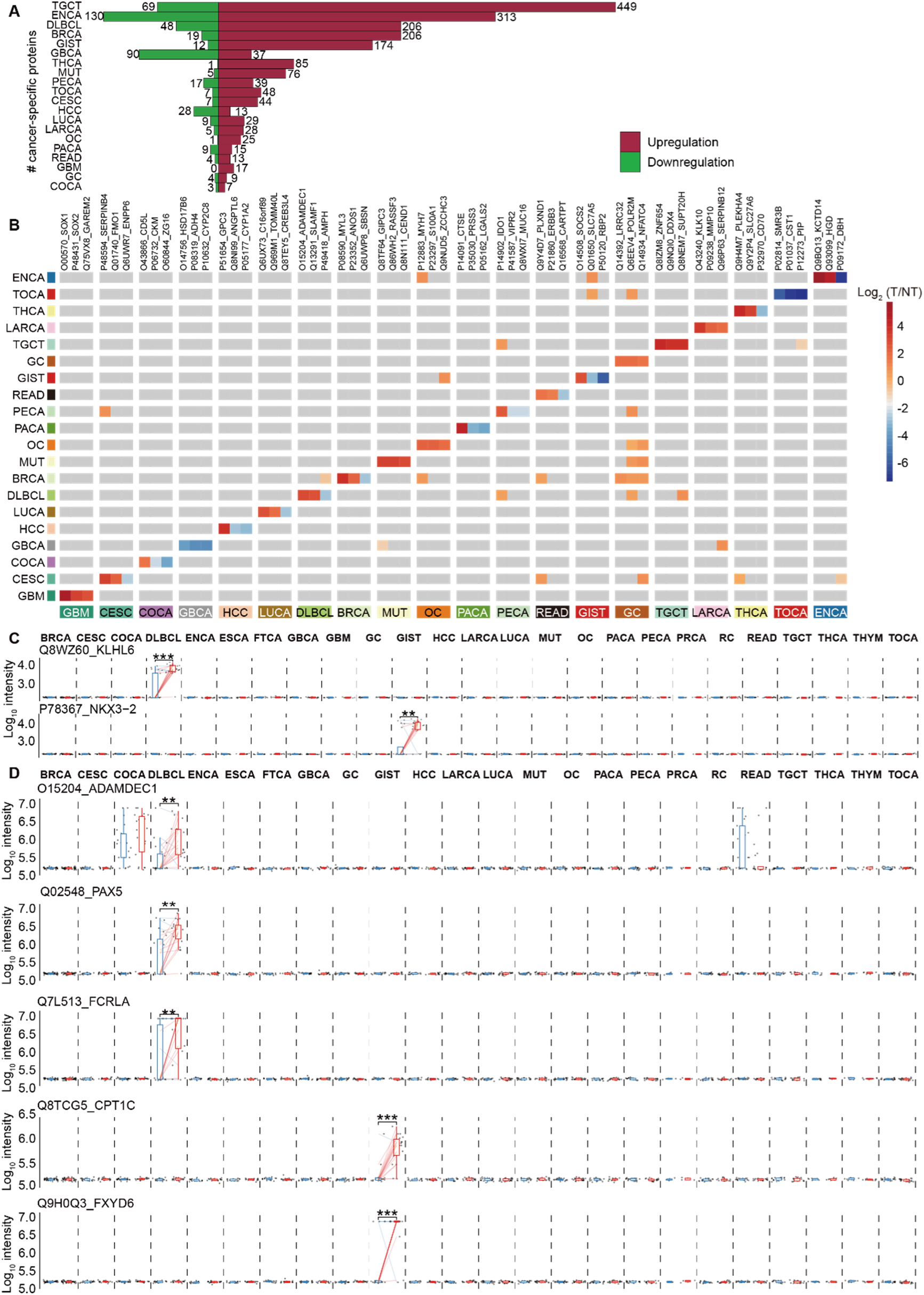
Cancer specific proteins in the pan-cancer dataset. **A,** Number of cancer specific proteins. **B,** Cancer specific proteins. Only 60 proteins are shown with the top 3 highest fold-change of each cancer type. **C,** PRM validated cancer specific proteins. Box plot shows protein expression; center lines indicate the medians and center points indicate the means; box limits represent the first and third quartiles; whiskers extend 1.5 times the interquartile range from the quartiles to outliers; translucent scatters show all sample points. Colored lines connect the carcinoma-adjacent pairs, while red lines mean up-regulations and green lines mean down-regulations. BH-adjusted *P* values are indicated by stars, * p<0.05, ** p<0.01, *** p<0.001.

**Figure S7.**
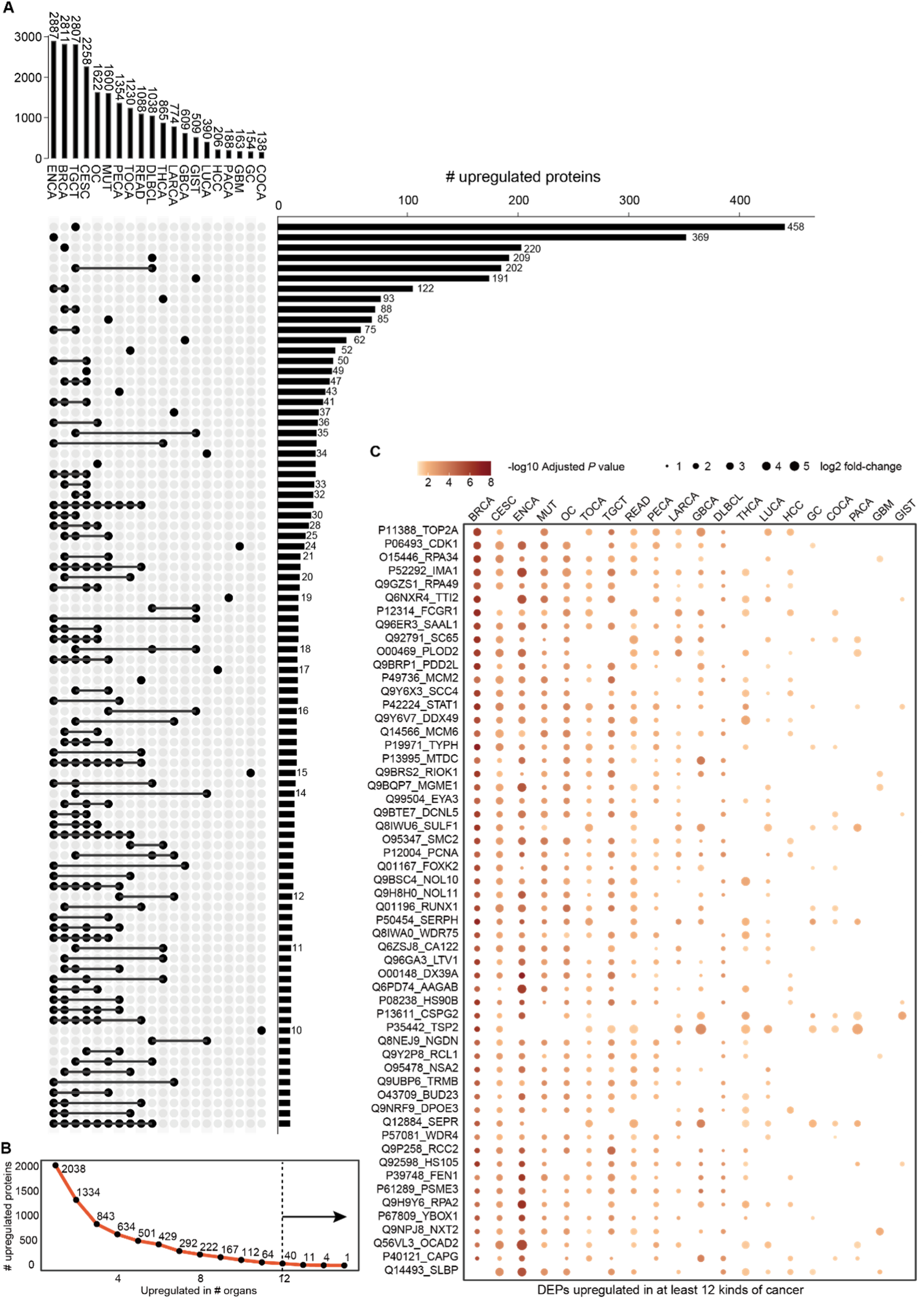
Most Common upregulated proteins among tumors. **A,** Sets are arranged in descending order according to the number of proteins they contain. Sets with the same number are only marked once. Only sets with at least 10 proteins are shown. **B**, Number of upregulated proteins in each group, with groups defined by the number of organs showing protein upregulation. **C**, Common differential expression illustrated by analyzing proteins upregulated in at least 12 organs.

