## Supplementary information for "Spatial distribution of the proteome in human body and cancers"

### Supplementary results

#### Anatomically dissectible resolution of human sample collection

In the sample repository, we collected histological healthy tissues mainly from a male and a female participant (**Extended data figure 1A, Figure E1A** for short) in an anatomy resolution. In total, we have 58 roughly partitioned tissue types, among which 24 were newly added compared with HPA[1] and TSomics[2] (**Figure E1A** ); and meanwhile 251 detail-partitioned tissue types, among which 212 were newly added (**Figure E1A** ). For in-situ tumor and its paired non-tumor samples, we recruited 6-52 patients for each of the 25 cancer types (**Table S1**). To handle the various tissue types in our collection, we applied the pressure cycling technology (PCT)-assisted sample preparation[3,4] along with in solution digestion and in-gel digestion method to extract proteins and digested them into peptides for the following DDA and DIA analysis.

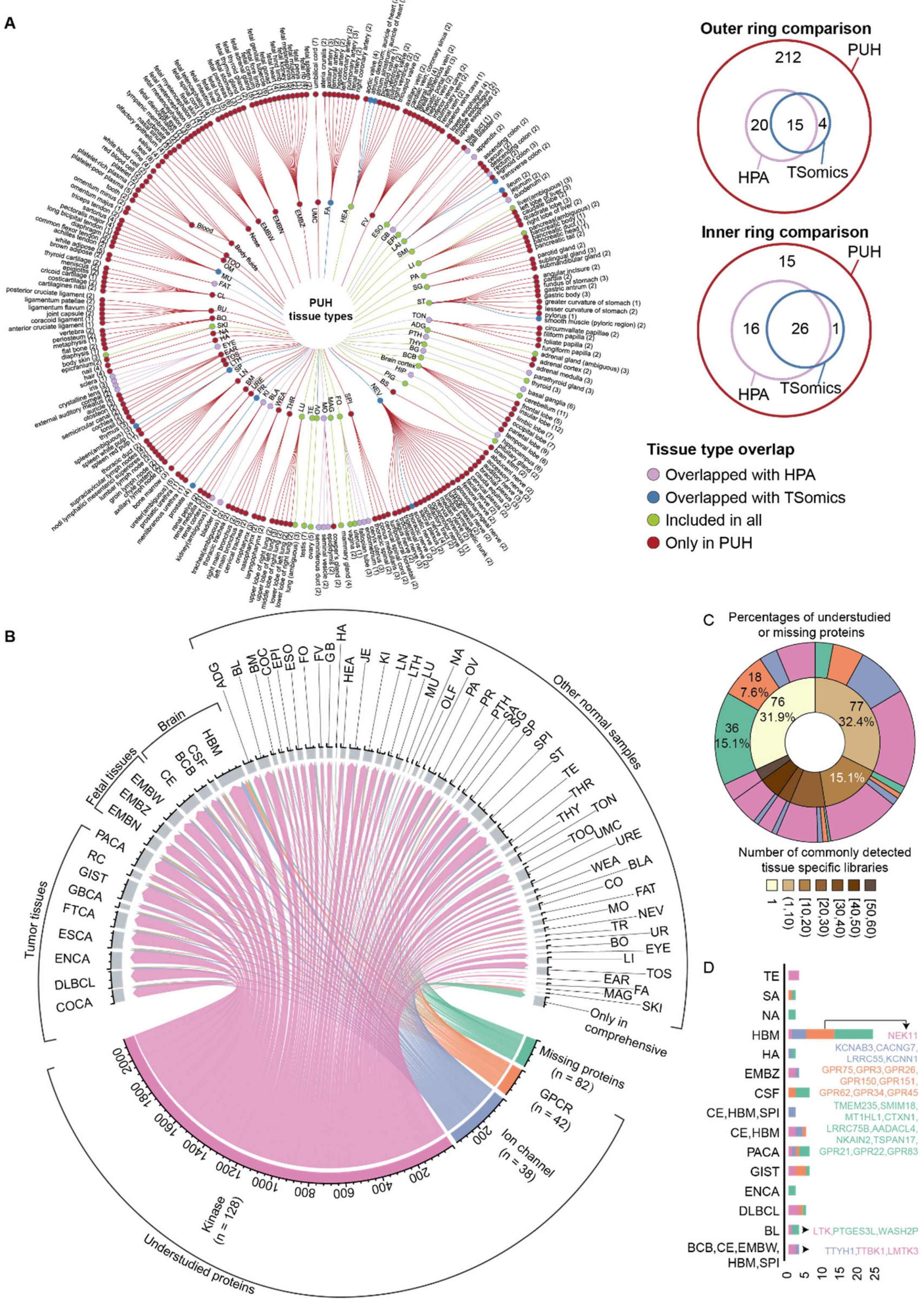


**Extended data figure 1. Overview of sample and library identification. A.** Tissue type overlap between this study and other published datasets. **B.** Number of identified understudied kinase, ion channel and G protein-coupled receptors (GPCR) and missing proteins in ddaPASEF data. **C**. Pie plot showing the distribution of the proportion of missing and understudied proteins commonly detected across multiple tissues. **D**. Stacked bar plot showing tissue types where the missing and understudied proteins were detected

### Construction of the comprehensive spectral library

To comprehensively quantify the proteome of human samples, we firstly built a "dictionary" ─ the MS spectral library on timsTOF Pro MS using DDA method, by analyzing pre-fractionated peptide sample pools per tissue type (**Figure S1, Table S2**). Altogether, we analyzed 1028 DDA files from 65 types of human specimens and generated 65 tissue-specific sub-libraries using FragPipe[5-8]. As shown in Figure S3B, the testis tissue-specific library contains the highest number of proteins (8929 proteins), while the tooth library has the fewest (1316 proteins). Evaluation of the quality of the comprehensive library was performed based on the statistical and plotting functions of DIALib-QC[9], with optimizations and reimplementations in R, incorporating protein coverage and missed cleavage distribution, thereby demonstrating the robustness of the library for downstream analyses (**Figure S1B-L**). By leveraging all the DDA files, the generated library contains 22,539,157 peptide spectrum matches (PSMs), leading to the identification of 15,332 proteins from 484,615 peptides. To ensure the quality of the spectral library, we evaluated the characteristics of the comprehensive library as shown in Figure S1B to S1L. The majority of precursor m/z values were settled between 300-1000 Th, with +2 and +3 precursors accounting for over 90% of the total. Meanwhile, these precursors exhibited a strong correlation in their chromatographic characteristics. Almost all the proteins have over six razor or unique peptides and almost every precursor has over six fragments. Most proteins in the dataset have protein coverage exceeding 20%. Our results showed a higher median protein coverage compared to the proteome map (33.43% versus 28%).

We also compared the comprehensive library with two of the largest human peptide-based resources, PeptideAtlas[10] and ProteomeDB[11]. PeptideAtlas contains curated peptide information collected from the proteomics community over the past decade while ProteomeDB achieved 92% proteome coverage by integrating about 17,000 human proteomic profiles. Notably, almost half of the peptides we identified were not deposited in ProteomeDB and still discovered over 70,000 novel peptides and 449 novel proteins not recorded either in the PeptideAtlas or ProteomeDB (**Figure S1M and S1N**). It is reasonable to attribute the expansion in protein identification to the breadth and depth of our proteomic analysis because we expanded the diversity of tissue types compared to previous studies **(Figure S1M and S1N**).

The newly built-up spectral library and the deep analysis enabled us to identify 82 missing proteins from Human Proteome Project as proteins lacking evidence at the protein-level (**Figure E1B**). Most of the missing proteins detected were distributed on chromosome 1 and chromosome 11 (**Figure E2A**), which was consistent with the HPP report[12], while they were detected in our comprehensive spectral library (**Figure E2B**). We validated the presence of the protein Pannexin 3 (PANX3) in cochlear samples, although it used to be detected only in other mammals (mice, rats) [13,14]. Our data further suggested that PANX3 may also play a role in human hearing. Besides, the existence of two orphan G protein-coupled receptors (GPCRs) were validated in human brain tissues (**Figure E2C**).

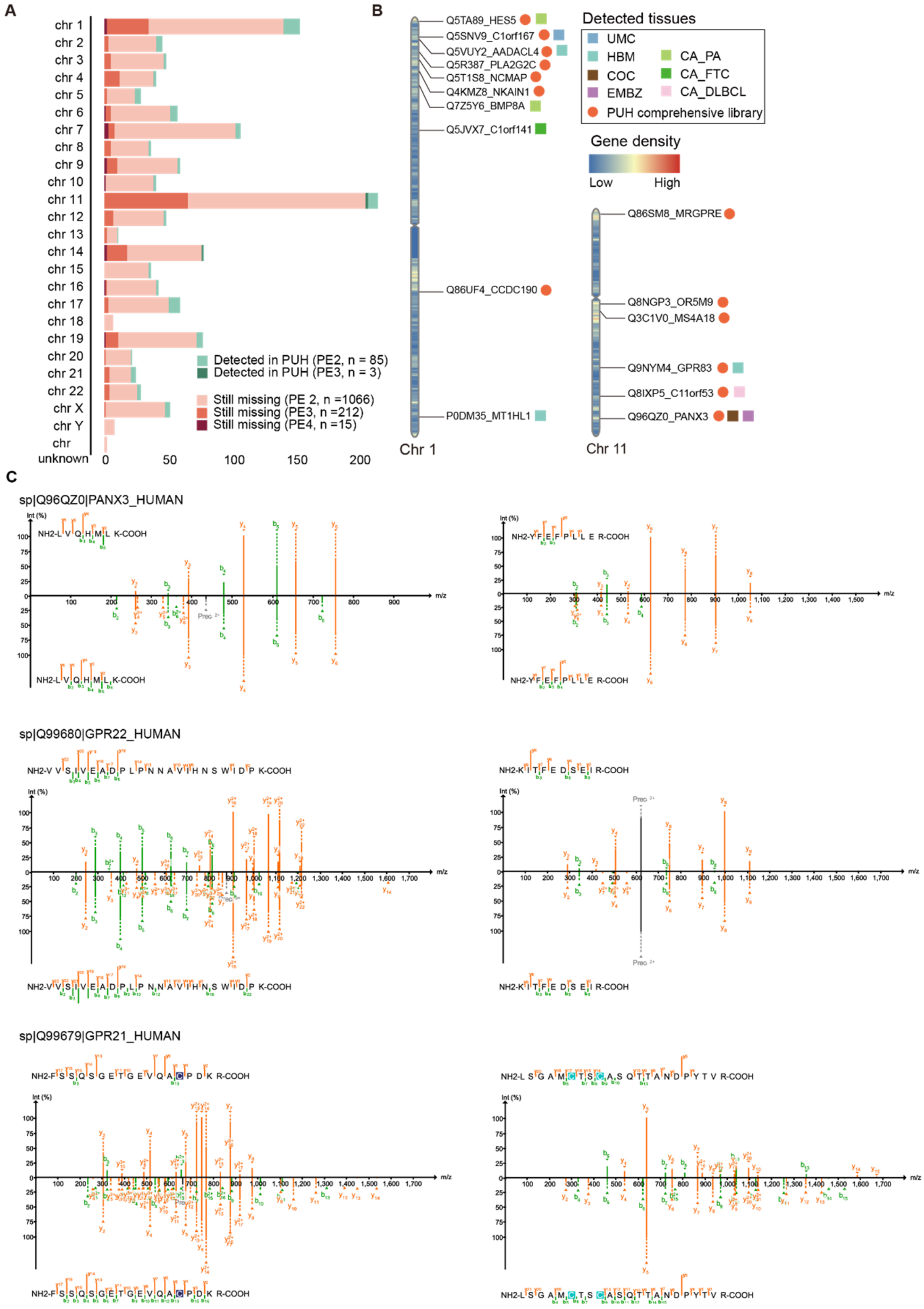


**Extended data figure 2. Missing proteins in comprehensive and tissue specific spectral libraries. A**. Bar plots showing the number of detected and undetected missing proteins on each chromosome in our data. **B**. Missing proteins on chromosome 1 and 11. Shapes represent the detected tissues of missing proteins. **C**. Annotated fragment ion spectra for proteotypic peptides of missing proteins in human tissue samples (upper) and pooled synthetic peptide sample (bottom).

### Data quality of the quantitative proteomic analysis

In summary, we used PCT to ensure the consistency in sample preparation for most sample types except for urine, saliva, and plasma, which were prepared by two-step overnight enzyme digestion method (**Figure S2A**). During the sample preparation process, a subset of 1579 samples was randomly selected as biological replicates, resulting in a total of 1635 peptide samples for DIA analysis. During the data acquisition process, a subset of samples was randomly selected for duplicate injections, resulting in a total of 1781 human sample DIA files. In total, 1900 DIA mass spectrometry files were generated. We used the comprehensive spectral library and DIA-NN to analyze these 1900 raw DIA files, resulting in a quantitative matrix of 13,477 proteins (**Figure S2A**).

Protein and peptide identification showed significant differences in different tissue types (**Figure S2B**). The numbers of proteins and peptide segments also varied within the same tissue type, possibly due to the structural and functional heterogeneity within the tissue type and the presence of cancer involvement. Generally, the number of protein identifications in Tumor (T) and Non-tumor (NT) samples is higher than that in Normal (N) samples within the same tissue type (**Figure S2B**). Among all samples, the maximum protein identification of a single-shot DIA was found in a colorectal cancer sample (9505 proteins), while the minimum protein identifications were found in plasma (a mean of 715 ± 134 proteins, n=13) and lens samples (777 and 875 proteins, n = 2).

Next, we performed quality control (QC) analysis on the quantification results of the obtained protein matrix. We found that most coefficient of variations (CVs) of the protein abundances in the 117 pooled peptide samples and 100 groups of replicates were low, with median values of 0.17 and 0.10, respectively (**Figure S2C**). Most of the correlation coefficients of protein abundances among the pooled samples and the replicates were high, with a median Pearson's correlation coefficient of 0.96 and 0.97, respectively (**Figure S2D**). By exploring the two-dimensional distribution of the samples, we evaluated whether the quantification results were affected by batch effects. Through principal component analysis (PCA) of the proteome, we found that the sample clustering was more influenced by sample types rather than batch factors such as acquisition month (**Figure S2E and S2F**).

For pan-cancer analysis, we filtrated a total of 1,170 paired tumor and non-tumor clinical samples, as well as companion pooled peptide samples (**Figure S3A**). Subsequently, we removed proteins that were completely missing in these samples, resulting in a matrix containing 1170 samples and 13,396 proteins. We conducted quality control analysis on this pan-cancer protein matrix to evaluate the reproducibility and batch effect in the pan-cancer proteomic matrix. We found that most CVs of the protein abundances in the 61 pooled peptide samples and 52 groups of replicates were low, with median values of 0.19 and 0.10, respectively (**Figure S3B**). Most of the correlation coefficients of protein abundances among pooled peptide samples and 52 groups of replicates were high, with a median Pearson's correlation coefficient of 0.97 for both (**Figure S3C**). Batch effect of pan-cancer samples was evaluated by PCA (**Figure S3D and S3E**). The analysis did not identify any sample clusters based on batch factors.

In summary, QC analysis indicated excellent reproducibility and minimal batch effects within the pan-cancer dataset, providing assurance for subsequent data analysis.

### Identifying enriched proteins for benign tissues

To clarify the proteomic characteristics of different human tissues, we analyzed the relative protein intensities among roughly partitioned tissue types of normal samples by post-mortem anatomy, except for the body fluids and embryo samples due to the huge differences of proteomes (**Figure S2G**). A total of 497 samples from 51 rough tissue types from 9 patients were selected. Firstly, we visualized the similarity of proteome between tissue types. From h-clust tree (**Figure 3A**), we found that physiologically related samples, like cerebral cortex and spinal cord, were more likely to cluster together. Few exceptions include: 1) peripheral nerve (NEV) and fat; 3) uterus and ureter; 4) throat and salivary gland. We found samples of these tissue types are dispersedly distributed in the t-SNE map (**Figure S2G**), and similarity of one sample would cause the similarity of the whole tissue type on one-dimension h-clustering, as most tissue types have few samples.

To identify the enriched proteins for each tissue, we classified 12,392 proteins in 51 tissue types into six groups according to the HPA criteria[15]: not detected, tissue enriched, group enriched, expressed in all, tissue enhanced, and mixed, with the composition of these categories shown in Figure S4A and Table S3. The proteins expressed in all tissues constituted around a quarter in the quantified protein IDs, while around three quarters of the total abundance (**Figure 3A**). Brain possessed the most tissue enriched proteins, whereas the summed abundance ratio of tissue enriched proteins to all identified proteins was highest in liver (**Figure 3A**). The formal has also been reported in previous studies due to the distinct biological function of brain, whereas the latter has been ignored. As shown in Figure S4B and S4C, liver enriched proteins ranked in the top 20% of the liver proteome, while brain enriched proteins mostly ranked in the bottom 80% of the brain proteome. We inferred that the high abundance ratio of tissue enriched proteins might be related to the higher burden of metabolism as indicated by the dominant metabolic pathways enriched by liver enriched proteins (**Figure S4D**). Most of the brain tissues were enriched in neural-related pathways (**Figure S4E**), suggesting that the enriched proteins could reflect the primary biological functions of the tissues. Collectively, we have developed a more comprehensive view of the proteomic landscape of different human tissues, highlighting their tissue-specific expression of proteins. These findings could have important implications for understanding the molecular mechanisms underlying tissue-specific functions and diseases.

### Dysregulated proteins in other pan-cancer datasets

We performed a thorough comparison of DEPs with the TCGA and CPTAC datasets to identify robust DEPs among different datasets (**Figure S5A, Figure S5B**). Regardless of the variances in cohorts and expression profiling methods, we still observed much consistency among these pan-cancer databases. We found 15,806 dysregulations were agreed upon for cancer types in our data and TCGA datasets (**Figure S5A, Table S6**), comprising 14,466 upregulations (5978 proteins) and 1340 downregulations (783 proteins). Similarly, 28,266 matched dysregulations on 8361 DEPs of 16 cancer types were identified between our data and the CPTAC dataset. Of these dysregulations, 9026 DEPs were shared in both datasets in the same cancer types (**Figure S5B**), including 7984 upregulations (3702 proteins) and 1042 downregulations (556 proteins). Specifically, 5068 upregulations (2878 proteins) and 470 downregulations (303 proteins) were consistently identified as significantly upregulated or downregulated proteins among three datasets from COCA, HCC, LUCA, GC, and LARCA.

We also performed a comprehensive analysis by cross-referencing cancer dysregulated proteins with both favorable and unfavorable prognostic datasets obtained from the Human Pathology Atlas[16]. We identified 6164 overlapping proteins from 11 types of cancer (**Figure S5C, Table S6**), exhibiting potential as biomarkers and therapeutic targets for the treatment of these cancers.

**Cancer specific DEPs associated with both tissue differentiation and oncogenesis**

Among GBM specific DEPs, SOX1, SOX2 and GAREM2 had the highest fold changes between T and N (**Figure S6B**). In GBM, developmental fate decisions are dictated by master transcription factors (TFs). A subset of stem-like tumor-propagating cells (TPCs) appeared to drive tumor progression and underlie therapeutic resistance. SOX2 was found to be one of a core set of neurodevelopmental TFs involved in binding and activating TPC-specific regulatory elements, and reprogramming differentiated GBM cells to “induced” TPCs. Additionally, SOX1 and SOX2 were found to individually reprogram GBM cells to enhance spherogenesis[17]. GAREM2 has been reported as a brain-specific Grb2-associated regulator of extracellular signal-regulated kinase (Erk)/mitogen-activated protein kinase (MAPK) (GAREM) subtype, contributing to neurite outgrowth of neuroblastoma cells by regulating Erk signaling[18]. Kelch-like protein 6 was selected as a potential DLBCL-specific biomarker in our dataset (**Figure S6C**), and it has been proved to be relevant to the differentiation of B cells in mouse models[19] and a tumor suppressor gene for the DLBCL. The loss of KLHL6 favored the growth and survival of DLBCL both in vitro and in xenograft models[20]. We identified NKX3-2 as a cancer-specific biomarker for GIST (**Figure S6C**). NKX3-2 is a transcriptional repressor that acts as a negative regulator of chondrocyte maturation and is an important regulator of gastroduodenal tract morphogenesis. Downregulation of NKX3-2 indicates poor prognosis for gastric cancer by promoting tumor migration and invasion via TGF-β-induced epithelial-mesenchymal transition[21].

### Dysregulated proteins as the drug targets in clinical trials

To discover potential cancer therapies under investigation, we searched 1266 pairs of cancer indications - drug entries constituted by the 289 drug candidates and their targeted cancers on the ClinicalTrials.gov database (version as of June 2, 2023). We obtained 2089 clinical trials related to 40 drugs and 81 dysregulated proteins in 18 cancers with the last update posted from 2005 to 2023 (**Figure 4B**). The drugs were classified into the following kinds according to their mechanism: RTK inhibitor (81.71%), immunotherapy (14.21%), antibody-drug conjugate (1.91%), hormonal agent (1.29%), antibody (0.57%), and angiogenesis inhibitor (0.29%). Excluding 203 trials with unavailable phases, 16.38% (309) are in phase 1 or early phase 1, 63.84% (1204) are in phase 2, and 18.03% (340) are in phase 3. Only 1.75% (33) of the trials were in phase 4 and matched to 9 drugs, twelve of which were related to the use of Bevacizumab as a vascular endothelial growth factor A (VEGFA) inhibitor for the treatment of OC and ENCA[22]. Bevacizumab was also used for non-small cell lung cancer, colorectal cancer, renal cell carcinoma, and cervical cancer supported by phase 3 clinical trials, but these indications were not mapped because VEGFA was not identified as being dysregulated in these types. Besides, few ADC drugs were found. Enfortumab vedotin, an ADC drug targeting Nectin-4, is used in combination with pembrolizumab to treat advanced urothelial cancer in a phase 3 clinical trial[23]. Although not yet approved by the FDA, Enfortumab vedotin is expected to be approved in subsequent trials. More annotations of clinical trials for DEPs could be found in Table S6. Beyond these protein targets mapped with DrugBank and clinical trials, we believe that this database would further facilitate the development of new therapies or biomarkers, as targeting dysregulated proteins or related pathways has been frequently applied in biomedical research.

### Targeting the locally enriched dysregulated proteins

We found a significant difference in the number of LEDEPs among different tissue types (**Table S9**). Overall, the numbers of upregulated and downregulated LEDEPs were balanced, while LEDEPs in a certain tissue type were tended to be either upregulated or downregulated (**Table S9**). Among all the tissues, liver had the highest number of LEDEPs, including 22 downregulated enzymes and one downregulated transporter (**Table S9**). We performed Ingenuity Pathway Analysis (IPA) for the biological functions of these 23 proteins and found that most of them were involved in endogenous and exogenous metabolic pathways (**Figure 6, Table S9**). The reduction of these proteins suggests a decrease in the proportion of normal differentiated liver cells or impaired metabolic function in cancer lesions. We found that ALDOB protein in the glycolysis pathway was downregulated in HCC, which has also been reported in previous literature[24]. ALDOB has been reported to be significantly associated with the prognosis of HCC and is a candidate prognostic biomarker[25]. Furthermore, GYS2 protein, a key enzyme in glycogen synthesis pathway, and SDS protein, which is important in fatty acid metabolism, were also liver LEDEPs. Metabolic alterations of glycogen, amino acid and fatty acid have been widely reported in HCC[26], providing biological insights and therapeutic opportunities for HCC[27]. Bai et al. applied the expression levels of aspartate metabolic pathways to divide clinical samples of HCC into two molecular subtypes: Group 1 defined as a high aspartame metabolism subgroup, characterized by higher expression of asparagine synthesis gene ASNS compared to Group 2 (low aspartame metabolism group), and lower expression of asparagine degradation gene ASPG, which is one of the liver LEDEPs. They found that Group 1 had a worse prognosis[28]. One of the liber LEDEP, GLS2, is a key regulatory factor in glutamine decomposition and has been found to have tumor suppressor functions[29]. More liver LEDEPs related to amino acid metabolism can be seen in Table S9 and Figure 7. It is worth mentioning that CYP1A2, CYP1B6, SULT1A1, and SULT1A2 were significantly enriched in the metabolism pathway of xenobiotics, including the degradation of substances like melatonin, dopamine, amphetamine, and nicotine (**Table S9**). The downregulation of these proteins in cancer suggests a weakening of drug metabolic functions in HCC. In addition to the above-mentioned metabolism-related enzyme proteins, we also found that transport protein ABCC1 is a liver LEDEP. ABCC1 is an ATP-dependent membrane-bound transporter. It is related to drug resistance and malignant potential of tumors and has been reported as a potential therapeutic target and prognostic-related protein in HCC[30].

We observed 11 upregulated and one downregulated LEDEPs in OC, including nine nuclear proteins and five transcription factors including WT1, the well-known OC biomarker. Wilms tumor protein 1 (WT1) is a well-known prognostic and diagnostic biomarker for OC. Zhang et al. found that WT1 expression was significantly higher in ovarian cancer tissue compared to normal ovarian tissue, and knocking down WT1 in ovarian cancer cells inhibited cell proliferation, migration, and invasion. It was speculated that WT1 might promote ovarian cancer progression by activating the Wnt/β-catenin pathway[31]. Functional annotation of these 12 proteins using IPA indicated direct or indirect regulatory relationships between several transcription factors and STAT3 (**Figure 6**). STAT3 not only suppresses anti-tumor immune responses but also enhances cancer cell proliferation, migration, and survival. Our findings suggest that inhibiting STAT3 may be effective in ovarian cancer treatment, and we have also identified reports on the use of upstream receptor inhibitors for STAT3 in ovarian cancer[32] .

We identified seven upregulated LEDEPs in TGCT, including five nuclear proteins and two transcription factors. Functional analysis by IPA indicated enrichment of upstream regulatory factors STAT6, which regulates PMS1 and GTSE1. Literature research has shown that the STAT6 signaling pathway is highly activated in tumors and has been implicated in promoting tumor metastasis in colorectal cancer, non-small cell lung cancer, and melanoma[33]. Our data further highlighted the potential clinical value of STAT6 in TGCT. Notably, the upregulation of PMS1 was also observed in the nonobstructive azoospermia and mutations in the PMS1 gene are associated with an increased risk of developing testicular cancer[34].

According to our data, Napsin-A (NAPSA) is primarily expressed in the lungs and kidneys, and it is overexpressed in lung cancer samples, consistent with previous research finding[35]. NAPSA is important for maintaining lung alveolar tension and synthesizing angiotensin II in the kidneys. It is also a promising research biomarker for the diagnosis and prognosis of lung and kidney cancer. Another LEDEP in the lungs is Collagen type VI alpha 6 chain (COL6A6), which is downregulated in lung cancer, consistent with the literature[36]. COL6A6 belongs to the type VI collagen family and is encoded by a coding region of 6,789 base pairs. It plays an important role in maintaining cellular structural integrity and regulating cellular functions as an extracellular matrix protein. It can also be measured in the blood by MS[10], which makes it a potential blood marker for lung cancer.

### Materials and Methods

#### Collection of tissue samples

The autopsy samples involved in this study were collected from the body donation center of Dalian Medical University. Postmodern interval was within 10 hours. The project has been approved by the Ethics Committee of Dalian Medical University.

All the carcinoma and its paired adjacent samples from surgery were collected from Harbin Medical University Cancer Hospital. The project has been approved by the Ethics Committee of Harbin Medical University Cancer Hospital. Written informed consent for each patient was collected. The samples mentioned above are stored in formalin foxed paraffin embedded format.

Autopsy specimens of fetus were collected from Shenzhen Baoan District Maternal and Child Health Hospital (during 20200616 to 20200916) died of abortion. Postmodern interval was within 10 hours. The project has been approved by Shenzhen Baoan District Maternal and Child Health Hospital. Written informed consent for each patient was collected. The fetal samples are stored in formalin in 4°C before use.

### Autopsies and tumor tissue sample preparation for LC-MS analysis

The sample preparation process is much the same as described in the previous article [3,37]. In short, we punched each FFPE sample and weighed about 1.0 mg. Heptane was used for dewaxing, gradient (100%, 90%, 75%) ethanol for hydration, and 0.1% formic acid for acid hydrolysis. The obtained product was then placed in PCT microtubes, and Tris-HCl (pH=10, freshly prepared) was added at 95°C for 30 mins for alkaline hydrolysis. After rapid cooling, lysis buffer (6M uera, 2M thiourea), Tris(2-carboxyethyl)phosphine (TCEP) and iodoacetamide (IAA) were added for reductive alkylation. PCT-assisted lysis was performed in the Barocycler model NEP2320-Enhanced (Pressure BioSciences, Inc, South Easton, MA) at 45,000 psi, 30 s HP, 10 s AP and 30℃ for 90 cycles. LysC (enzymatic substrate concentration ratio 1:80, mass spectrometry, Wako, Richmond, VA, USA) and trypsin (enzymatic substrate concentration ratio 1:20, sequencing grade modification, Promega, Madison, WI, USA) were then added for digestion. The process was conducted with the assistance of the PCT under conditions of 20,000 psi, 50s HP, 10s AP, and 30℃ for 120 cycles. After the enzymatic hydrolysis process was terminated by 10% formic acid, we used the C18 cartridges (Waters, Milford, USA) for desalination.

The dried samples were resuspended with buffer A (HPLC grade water, pH=10). The fractionation was performed using a Thermo Fisher Ultimate 3000 RSLCnano System (Thermo Fisher Scientific™, San Jose, USA) with an HPLC C18 column (diameter 4.6 mm, length 25 cm, pore size xx, particle size xx). The flow rate was 0.5μL/min. The gradient was 5% -95% buffer B (98% acetonitrile, pH = 10) in 60 mins. A total of 60 fractions were collected. Then 60 fractionations were combined into 10. The combined samples were evaporated using vacuum centrifugation (CentriVap, Labconco, Kansas City, USA) at 45℃.

For bone tissue, we cut a piece of them, approximately one-fifth the volume of a 2 mL centrifuge tube. The teeth were shattered with a cleaned hammer and placed into the 2 mL tube. After the dewaxing and alkaline hydrolysis described above, we added lysis buffer and grinding beads (about one-third the volume of the 2 mL tube). The sample was frozen in liquid nitrogen, ground in a tissue grinder for one minute, and refrozen in liquid nitrogen. This procedure was repeated 3 to 5 times until the sample turned into a bone slurry (except for the teeth). We then added 1 mL of 10% formic acid solution and incubated the sample at 4°C overnight for digestion. Afterward, approximately 4 mg of the insoluble wet weight was transferred to a PCT tube, and 150 µL of Tris-HCl (pH=10) was added to neutralize any excess acid, followed by removal of the supernatant. Then, 20 µL of Tris-HCl (pH=10) and 30 µL of 6 M urea/2 M thiourea lysis buffer were added, and subsequent PCT-assisted lysis and enzymatic digestion were performed as described above.

Hair samples (4 cm) or nail samples (1 mg) were immersed in 50% methanol or ethanol, chopped, and vortexed, then transferred to a PCT tube. To the tube, 30 µL of lysis buffer (30% trifluoroethanol, Tris-HCl buffer, pH=8) and 2.5 µL of 200 mM TCEP solution were added, and the tube was sealed with a PCT pestle and placed in a PCT device. The samples underwent 180 cycles in the Barocycler (50 s at 45,000 psi, 10 s at atmospheric pressure, 70°C). After cooling to room temperature, 2.5 µL of 800 mM IAA solution was added, and the samples were incubated in the dark at room temperature for 30 minutes with gentle mixing (800 rpm). After dilution with 150 µL of Tris-HCl buffer (pH=8), 1.25 µg of Lys-C protease and 5 µg of trypsin were added, and the tube was sealed and subjected to 120 cycles in the PCT (20,000 psi for 50 s, atmospheric pressure for 10 s, 30°C). After the cycles, 15 μL of 10% TFA solution was added to stop the digestion, and the samples were desalted using C18 tips.

### Sample collection and preparation for blood, saliva, urine, hair and nail

The whole blood was harvested by venipuncture of participants into commercial EDTA-containing sampling containers. And we can separate it into PRP1, erythrocyte, platelet-rich plasma (PRP), platelet-free plasma (PPP), the pure platelet fraction referring to the article published in 2019[38]. PRP1 is the supernatant in the first centrifuge at 200 g for 10 min. Then we divide 1ml whole blood from the anticoagulation tube into a 50ml centrifuge tube, add 10ml 1X red blood cell lysate (Invitrogen, catalog number 00-4333), 10 minutes at room temperature (strictly controlled time), add 25ml PBS (total volume 36ml) to stop the reaction, then divide into 15ml centrifuge tubes (9ml each tube), centrifuge at 500g for 15 minutes at 4°C. Aspirate and discard the supernatant. At this time, allow 200ul of liquid to remain in each 15ml centrifuge tube. Pipette the remaining liquid and cell pellet from each tube into a 1.5ml centrifuge tube. Centrifuge at 500g for 5 minutes at 4°C, and then aspirate the supernatant as much as possible. Leaving the cell pellet. Wash the cell pellet with 1*PBS three times and centrifuge to get a clean white blood cell pellet (500g for 5 minutes, 4°C).

The first part is the depletion of top 14 proteins in blood, PPP, PRP, and PRP_1: samples taken out from -80 °C were immediately put on ice. Equilibrate the depletion column to room temperature. Following the manufacturer's instructions (Thermo Scientific, High-Select™ Top14 Abundant Protein Depletion Resin, cat # A36371).

The second part is protein concentration which is following the manufacturer's instructions (Thermo Scientific, Pierce™ Protein Concentrators PES, 3K MWCO, 0.5 mL, Cat #88512). Then add 500ul 8M urea/100mM ammonium bicarbonate (AAB) to the concentrator and spin at 12000 × g for 40 min. Finally, adjust the remain volume to 50ul.

The third part is protein digestion we could refer to the published article[39]. However, the times of TCEP and IAA process were both 40min. Then the protein extracts were digested with lysC and trypsin to the sample at 1:50. Digested peptides were cleaned using C18 (Thermo Fisher Scientific, cat # 60209-001).

The fourth part is peptide fractionation as we have described in Nie et al.[40]. Briefly, peptides were separated into 122 fractions, which were consolidated into 10 fractions. The samples were separated at a flow rate of 1 mL/min using a gradient from 5% acetonitrile (ACN) in 10 mM ammonia (pH = 10.0) to 8% in 7.5min, the percentage increased to 18% in 30min and to 32% in 55min then to 95% in 61min, this 95% composition remained six minutes, finally it dropped to 5% in 68min. The fractions were combined as following strategy: 1) combine the 1st to 6th, 77th to 82nd and 97th, 98th, 107thand 108th fractions; 2) combine the 7th to 10th, 73rd to 76th, 93rd, 94th, 111stand 112nd fractions; 3) combine the 11st to 14th, 69th to 72nd, 89th, 90th, 115thand 116th fractions; 4) combine the 15th to 18th, 65th to 68th, 89th, 90th, 115thand 116th fractions; 5) combine the 19th to 22nd, 61st to 64th, 99th, 100th, 105thand 106th fractions; 6) combine the 23rd to 26th, 57th to 60th, 95th, 96th, 109thand 110th fractions; 7) combine the 27th to 30th, 53rd to 56th, 91st, 92nd, 113rdand 114th fractions; 8) combine the 31st to 34th, 49th to 52nd, 85th, 86th, 119thand 120th fractions; 9) combine the 35th to 38th, 45th to 48th, 101st, 102nd, 103rdand 104th fractions; 10) combine the 39th to 44th, 87th, 88th, 117thand 118th fractions. The combined fractions were subsequently dried and re-dissolved in 2% ACN/0.1%

formic acid (FA), and then analyzed by LC-MS (Bruker, timsTOF pro).

Around 2 mg samples were processed to generate peptide samples using accelerated PCT assisted sample preparation method as described previously (PMID: 32182071; PMID: 29071490). Briefly, the samples were lysis with 30ul buffer containing 8 M urea/2M thiourea (Sigma), reduction by Tris (2 carboxyethyl) phosphine (TCEP, Sigma) and alkylation by IAA in PCT (100cycles, 45kpsi, 30s HP and 10s AP at 30°C). Then the lysates were digested using PCT by a mix of Lys-C (enzyme-to-substrate ratio, 1:80) and trypsin (enzyme-to-substrate ratio, 1:20) (Hualishi Tech. Ltd, Beijing, China) in PCT (120cycles, 20kpsi, 50s HP and 10s AP at 30°C). The digestion was quenched by trifluoroacetic acid (TFA) (Thermo Fisher Scientific). Finally, peptides were cleaned by C18 spin columns (The Nest Group, Inc., MA), dried under vacuum and fractions were performed as above for MS analysis.

For the collection of saliva, we have strictly controlled the condition to minimize the pollution from food debris and oral bleeding according to the instructions from Salimetrics (https://salimetrics.com/saliva-collection-handbook/). In brief, we collected about 1mL saliva respectively from 4 healthy participants before eating meals and at least 45 min after brushing teeth with the records of the start and end time in a passive drool way. And we avoided such situation before collection: strenuous exercise; oral issues; intake of high sugar diet, highly sour diet, coffee, alcohol, nicotine and drugs. We collected the 1 mL of middle part of the morning urine. The saliva and urine samples were stored in -80°C before use. The following reduction, alkylation, digestion, and desalting steps were performed as described for plasma samples described in published article[39].

We collected the tail of hair and finger nail, then stored in 4℃ before use.

### LC-MS/MS analysis

The peptides re-dissolved in 98% water:2% ACN:0.1% formic acid (v/v) were separated and analyzed by coupled UHPLC-TIMS-MS/MS system. For each acquisition, the peptides were loaded onto Thermo Scientific™ Trap Cartridge (100 Å 5.0 µm, 0.3 mm × 5 mm), and separated by BPRC analytical column (120 Å 1.9 µm, 150 mm × 0.075 mm) with Bruker Daltonics™ nanoElute UHPLC system using a 95 min LC gradient at a flow rate of 0.3 µL/min. The composition of gradient was as follows: linearly increased from 2% B to 22% B in first 80 minutes, then from 22% B to 35% B in 10 minutes, and then from 35% B to 80% B in 2 minutes and maintained 80% B in the last 3 minutes. The mobile phase A was 99.9% water:0.1% formic acid (v/v) while mobile phase B was 99.9% acetonitrile:0.1% formic acid (v/v). And all reagents were MS grade.

Bruker Daltonics™ trapped ion mobility quadrupole time-of-flight mass spectrometer (timsTOF Pro) performed data acquisition. In both DDA and DIA mode, the MS full scans were acquired over an m/z range of 100 to 1700 and an ion mobility range of 0.60 to 1.60 $V \cdot s/cm^2$. The ion mobility over m/z heatmaps were filtered using an inclusion/exclusion polygon region with vertices (150, 0.6), (1300, 1.6), (1700, 0.6), and (1700, 1.6). In ddaPASEF mode, the method consisted of a MS full scan followed by 10 PASEF MS/MS scans, which consuming 1.17 s of total cycle time. The accumulation time and ramp time were both 100 ms. The target intensity was 20,000 and the intensity threshold was 2500. And in diaPASEF mode, the PASEF MS/MS scans were collected from 64 windows in an m/z range of 400 to 1200 and corresponding ion mobility range of 0.57 to 1.47 $V \cdot s/cm^2$ adapted from standard 16 diaPASEF scans scheme[41].

### Database searching and library generation

For generating consensus PASEF spectral libraries, we used FragPipe computational platform (https://fragpipe.nesvilab.org/) (version 18) with MSFragger[7,8] (version 3.5), Philosopher[42] (version 4.2.2) components and EasyPQP (https://github.com/grosenberger/easypqp/) (version 0.1.29) Python package. And our workflow was based on the FragPipe DIA_SpecLib_Quant workflow[5] after being modified accordingly.

The database searching of raw files was performed by MSFragger search engine, referring to a UniProtKB/Swiss-Prot Homo sapiens (UP000005640) database (containing 20,377 canonical protein sequences; downloaded on May 7th, 2020) appended with reverse protein sequences as decoys generated by the built-in Philosopher decoys generation tool. Then Philosopher was utilized for further evaluation, protein inference, and 2D FDR filtering with a threshold of 0.01, and built-in library generation script gen_con_spec_lib.py was utilized to build consensus PASEF spectral libraries with 0.01 FDR control on both peptide-level and protein-level. Most of the parameters were maintained as reported in the prior report [43], except for the application of semi-enzymatic tryptic digestion rule, 0.05 daltons of fragment mass tolerance, enabled razor algorithm and spiked iRT based linearly RT alignment.

### Spectral library evaluation

The information of precursor, fragment ion and modification containing in the spectral library were analyzed to evaluate its quality based on the algorithm modified from published DIALib-QC tool[9]. The Perl code for visualization was re-implemented in R. The ratios of sequence length, ion charge, peptide modification, fragment ion type, and missed cleavage were counted respectively and presented in bar charts. The distributions of precursor m/z and peptide sequence coverage were shown in histograms and the protein sequence database read-in was performed by seqinR package[44]. Pearson's correlation of iRT values between +2 and +3 ion charge states of the same peptide sequence was measured (R-squared) and the 2D kernel density estimation was performed using function stat_density2d in ggplot2 package.

We further compared the proteins and peptides in our spectral library with peptideAtlas (http://www.peptideatlas.org/builds/human) which consisted of 16,702 proteins and 2,701,150 peptides (by 2021.7.6), and ProteomeDB [https://www.proteomicsdb.org] which covered 13,553 proteins and 913,326 peptides (by 2021.7.6). The comparison results were shown in Venn diagram generated by R script.

### Analysis of DIA data by DIA-NN

DIA searching was performed by DIA-NN (version 1.8) with above FragPipe-generated spectral library with its existed protein inference results. As in the previous method[43], both MS1 and MS2 mass accuracy were set as 10 ppm, and Robust LC (high precision) quantification strategy and RT-dependent cross-run normalization were chosen for quantitative results. Moreover, the different runs were treated as unrelated and 0.01 FDR threshold on both precursor and protein levels was performed to generate the final reports with disabled MBR.

For shorter waiting time, the 1783 sample runs including replicates and 117 pooled sample runs were randomly grouped and searched on four different computers in parallel. And then, the 1900 runs were reanalyzed with existing .quant files to generate the final reports. The 23 mouse liver runs were searched with same settings as above for another quality control.

### Quality control and data preprocess

Before quality control analysis, the DIA matrix was removed proteins with a missing ratio of not less than 60%. The protein matrix was log2 transformed and filled missing values by zeroes to perform principal component analysis (PCA) for unsupervised estimation of batch effects of sample acquisition dates, instrument IDs, and timsTOF glass capillary changes. Moreover, the corresponding pooling data were extracted to calculate the coefficients of variation (CV) of protein abundances omitting missing values and Pearson correlation coefficients using all complete pairs of proteins on the samples. For both technical and biological replicates, protein abundance vectors were grouped by their unique sample ID and calculate the differences of quantified protein number, CV, and Pearson correlation coefficients using the same method as pooling data.

The protein matrix of carcinoma and adjacent data was quantile normalized of protein level intensities using the normalize.quantiles function provided by preprocessCore R package (version 1.58.0) [https://github.com/bmbolstad/preprocessCore] before log2 transformation and batch effect correction. Batch effects due to an over eight-month interval of sample preparation were corrected using the removeBatchEffect function provided by limma R package (version 3.52.4)[45]. Similar QC analysis was subsequently performed with the corrected matrix. What's more, we excluded body fluid, crystalline lens, hair, nail, and tooth from normal samples matrix before quantile normalization and log2 transformation. This matrix was used as normal samples matrix in subsequent analysis if not otherwise specified.

### Identification and quantification overview

The numbers of proteins in tissue specific libraries and DIA quantified peptides and proteins were visualized in circular bar charts using canvasXpress [https://CRAN.R-project.org/package=canvasXpress] R package and retouched using Adobe Illustrator. The anatomical classifications and sample types containing tumor, non-tumor, and normal were into consideration.

### t-SNE visualization

The whole matrix excluding pooled samples was standardized and dimensionality reduced using t-SNE with the Rtsne package in R. The perplexity parameter was set to (n-1)/3-1, where n was the number of samples. The two-dimensional result was visualized in a scatter plot.

### Differential expression of sample types

The differential expression from fetal, tumorous, paired non-tumorous to normal samples was assessed using three groups of two-side Wilcoxon signed-rank test and adjusting for multiple comparisons using the Benjamini-Hochberg procedure omitting missing values. Proteins with |fold change| of mean value not less than 1.2 and adjusted p-value < 0.05 were extracted as significantly differential expressed proteins. The upregulated proteins among three comparison groups as well as downregulated proteins were extracted respectively for GO enrichment using limma package in R. The GO biological process gene sets with genes between 10 and 5000 were selected and the p-value cutoff was set as 0.05. The top 10 significant pathways were shown in the bubble plot, with five pathways contributed by upregulated proteins and five pathways contributed by downregulated proteins.

### Tissue specificity analysis

The tissue specificity of quantified proteins of normal tissue samples with only autopsy specimens excluding body fluids, hair, nail, umbilical cord, and aborted fetus was estimated using modified HPA (Human Proteome Atlas) classification method [15]. The protein abundances of each tissue type were aggregated by median of all samples after imputing missing values to 0.5 times the minimal value in the whole matrix. We changed the categories of tissue enriched, group enriched, and tissue enhanced proteins from 5-fold higher abundance to 3-fold for our more varied tissue types.

### Pan-cancer analysis

The differential expression analysis was performed respectively by cancer types. For each kind of cancer type, a submatrix was extracted with tumorous and paired non-tumorous samples and removed proteins with a missing ratio of not less than 50%. Then the missing values were filled by a series of random numbers generated from a normal distribution with a mean of 1 and a variance of 0.001, which were then multiplied by 0.5 times the minimal value in the whole matrix. Two-side paired t-test and BH-adjustment of p-values were used to assess significant cancer dysregulated proteins. The adjusted p-value cutoff was 0.05 and only proteins with |fold change| of mean value not less than 2 were selected. Proteins dysregulated in only one kind of cancer were reported as cancer specific dysregulated proteins. The cancer dysregulated receptor tyrosine kinases (RTKs) were shown in heatmap after standardization and hierarchical cluster analysis with Euclidean distance. The cancer specific submatrices without NA imputation were combined to one for t-SNE visualization. And tissue specificity analysis was also performed to select cancer enriched proteins with the same methods mentioned above.

The cancer dysregulated proteins were filtered by TCGA curated pathways[46]. And then we counted the number of overlapped proteins for each cancer and each pathway to estimate the level of pathway dysregulation.

### Locally enriched proteins

The list of cancer specific dysregulated proteins was mapped to tissue enriched proteins from normal samples. Locally enriched proteins were defined as the intersection of them. The fold change cutoff of dysregulated proteins was set as 1.5 to extract more proteins for IPA analysis.

### Cross-referencing with external databases

The list of cancer dysregulated proteins was mapped to DrugBank online database[47] to find out potential targeted drugs. And the cancer-target-drug combinations were used to query clinical trials information from ClinicalTrials.gov online database [https://clinicaltrials.gov]. All matched clinical trials were visualized in a circular scatter plot.

We used the pan-cancer proteomic map (ProCan-DepMapSanger) drug response and CRISPR-Cas9 gene essentiality screens dataset[48] as a validation after filtering nc_fdr < 0.01, nc_beta < -0.1, and the protein-protein interaction between protein and drug target equals to "T". The gene names were transformed into Swiss-Prot protein IDs and mapped to our list of cancer dysregulated proteins. The beta without transcriptomics covariate, fold change of mean value, and cancer of each validated protein-drug relationship was shown in a scatter plot.

Genomics of Drug Sensitivity in Cancer (GDSC) dataset[49] was used as another validation. We filtered the result of ANOVA with COSMIC cell lines by FDR < 0.01 before mapping to our cancer upregulated proteins list.

The favorable prognostic, unfavorable prognostic and cancer enriched genes containing 14 kinds of human cancer were obtained from the pathology atlas dataset [https://www.proteinatlas.org/humanproteome/pathology], including head and neck cancer, thyroid cancer, lung cancer, liver cancer, testis cancer, prostate cancer, stomach cancer, colorectal cancer, breast cancer, endometrial cancer, ovarian cancer, cervical cancer, pancreatic cancer, and renal cancer. Our list of cancer dysregulated proteins was mapped to above public datasets respectively with the same cancer types as depicted by a Sankey diagram.

The list of cancer dysregulated proteins was also mapped to TCGA and CPTAC datasets by proteins and corresponding cancer types. And the results were shown in scatter plots.

### PRM validation

A total of 137 cancer specific dysregulated proteins were selected from pan-cancer data acquired before 2022 for PRM validation. Each selected precursor has no modification and no missed cleavage, with the peptide length ranging from 8 to 20 using Skyline[50] (version 21.1). In total, 56 peptide precursors from 47 proteins and 9 iRT peptide precursors were analyzed. Among the 47 cancer specific proteins, 14 were still identified as cancer specific. Skyline[50] was used to analyze and quantify the targeted PRM data before generating peptide matrix. And the peptide matrix was transformed into protein matrix using online ProteomeExpert Peptide2Protein tool[51] by the mean of the top3 precursor intensities and with log2 transformation and quantile normalization. The cancer enriched proteins and cancer specific dysregulated proteins were selected as the methods mentioned above after filling missing values with 0.1 times the minimal value in the whole matrix. And the validated proteins were shown in box plots.

**Validation of the detection of missing proteins through synthetic reference peptides**

Peptides were synthesized for PANX3 (SLAHTAAEYMLSDALLPDR, LVQHMLK and YFEFPLLER), GPR22 (VVSIVEADPLPNNAVIHNSWIDPK, KITFEDSEIR), GRP21 (FSSQSGETGEVQACPDKR and LSGAMCTSCASQTTANDPYTVR) by Jietai Biotechnology Co. (Nanjing, China) (http://www.synpeptide.cn/). Thees peptides were re-dissolved in 98% water:2% ACN:0.1% formic acid (v/v), mixed and analyzed by the same LC-MS method adopted in the library construction process.Tandem MS spectra of all candidate peptides were manually reviewed in Fragpipe.

Data analysis mentioned above were based on R (version 4.2.0).
